# An overview of the quality assurance and quality control of magnetic resonance imaging data for the Ontario Neurodegenerative Disease Research Initiative (ONDRI): pipeline development and neuroinformatics

**DOI:** 10.1101/2020.01.10.896415

**Authors:** Christopher J.M. Scott, Stephen R. Arnott, Aditi Chemparathy, Fan Dong, Igor Solovey, Tom Gee, Tanya Schmah, Sofia Chavez, Nancy Lobaugh, Nuwan Nanayakkara, Shuai Liang, Mojdeh Zamyadi, Miracle Ozzoude, Melissa F. Holmes, Gregory M. Szilagyi, Joel Ramirez, Sean Symons, Sandra E. Black, Robert Bartha, Stephen Strother, the ONDRI investigators

## Abstract

Large scale research studies combining magnetic resonance imaging data generated at multiple sites on multiple vendor platforms are becoming more commonplace. The Ontario Neurodegenerative Disease Research Initiative (ONDRI - http://ondri.ca/), a project funded by the Ontario Brain Institute (OBI), is a recently established province-wide natural history study, which has recruited more than 500 participants from neurodegenerative disease groups including amyotrophic lateral sclerosis, fronto-temporal dementia, Parkinson’s disease, Alzheimer’s disease, mild cognitive impairment, and cerebrovascular disease (previously referred to as the vascular cognitive impairment cohort). Because of its multi-site nature, all captured data must be standardized and meet minimum quality standards to reduce variability. The goal of the ONDRI imaging platform is to maximize data quality by implementing vendor-specific harmonized MR imaging protocols (consistent with the Canadian Dementia Imaging Protocol - http://www.cdip-pcid.ca/), monitoring protocol adherence, qualitatively assessing image quality, measuring signal-to-noise and contrast-to-noise, monitoring system stability, and applying corrections based on the analysis of images from two different phantoms regularly acquired at each site. To maximize image quality, this work describes the use of various automatic pipelines and manual assessment steps, integrated within an established informatics and databasing platform, the Stroke Patient Recovery Research Database (SPReD) built on the Extensible Neuroimaging Archive Toolkit (XNAT), and contained within the Brain-CODE (Centre for Ontario Data Exploration) framework. The purpose of the current paper is to describe the steps undertaken by ONDRI to achieve this high standard of data integrity. Data have been successfully collected for the past 4 years with the pipelines and assessments identifying deviations, allowing for timely interventions and assessment of image quality.

## 1. INTRODUCTION

Degenerative brain diseases are amongst the most menacing disorders looming in our aging society. These diseases typically impact key aspects of an individual’s identity such as personality, memories, or abilities, and are ultimately fatal. The direct and indirect costs of such diseases are immense. For example, 500,000 people are currently afflicted with Alzheimer’s disease and related dementias in Canada alone, with care costs estimated at 15 billion dollars annually. With the prevalence expected to more than double in the next 20 years, care costs will reach more than 153 billion annually (1).

The global landscape of brain research is constantly evolving, both to face new challenges and to keep pace with technological advancements. This “neuro-renaissance” has come about due to myriad factors such as recent advances in the development and validation of disease biomarkers, including those derived from magnetic resonance imaging techniques and newly developed PET ligands, and the prospect of impending financial and social burden.

Large multi-site studies with participant numbers often in the hundreds or thousands are becoming more common and present unique challenges, but also opportunities. Along with large numbers, many studies are also acquiring enormous amounts of data on each participant often including multiple imaging modalities such as MRI and PET, neuropsychometric testing, detailed genetics, gait and balance, eye tracking and imaging, and many others. For an overview of all ONDRI data acquisition platforms see Farhan et al. (2). The data acquired by such studies are now often referred to as “big data”, and with the power afforded by such detailed and thorough collection comes the mammoth task of ensuring consistency and compatibility of the data collected, as well as developing new techniques for analysis, where traditional methods now often prove inefficient and inadequate. Examples of these massive-scale studies include the Framingham Heart Study (3), the Three-City Study (4), and the Canadian Alliance for Healthy Hearts and Minds (http://cahhm.mcmaster.ca/). From an imaging perspective, the Alzheimer’s Disease Neuroimaging Initiative (ADNI) (5) was a pioneering multi-site imaging study that set the standard for longitudinal natural history imaging studies in neurodegeneration. Although many of these studies capture data from large numbers of individuals, often the depth of phenotyping or information captured is limited, with notable exceptions being the Human Connectome Project (6), and the recently launched UK BioBank study aiming to capture imaging and other data on 100,000 participants (7).

With the creation of the Ontario Brain Institute (OBI - http://www.braininstitute.ca/) in 2010 it was recognized that a robust databasing and storage system would be required to securely store and share a vast collection of varying data types including brain imaging, genetic analyses and proteomics data. In response, Brain-CODE (Centre for Ontario Data Exploration) was built in partnership with the Indoc Consortium (http://www.indocresearch.org). The Ontario Neurodegenerative Disease Research Initiative (ONDRI) project (2) (http://ondri.ca/) was launched shortly thereafter. Briefly, ONDRI focuses on five critical areas of degenerative brain disease including amyotrophic lateral sclerosis (ALS), fronto-temporal dementia (FTD), Parkinson’s disease (PD), cerebrovascular disease (CVD – previously referred to as the vascular cognitive impairment (VCI) cohort), and Alzheimer’s disease (AD) and amnestic mild cognitive impairment (MCI). Baseline recruitment of more than 500 participants occurred at 12 research institutes across Ontario with extensive testing at baseline, and repeated annual assessment is continuing for at least two additional years per participant. These ambitious evaluations include a broad battery of neuropsychological tests, assessments of gait and balance, genomics and proteomics, ocular (retinal) imaging, eye tracking, and neuroimaging, all of which undergo extensive neuroinformatics and biostatistical oversight in order to ensure data standardization, minimize data errors, and ultimately identify useful data trends (i.e., potential disease biomarkers). One of the unique aspects of this study is the depth of phenotyping of the individuals from each of the disease themes, allowing for comprehensive comparisons within and between diseases, both cross-sectionally and longitudinally as depicted in Figure 1.

**Figure 1:**
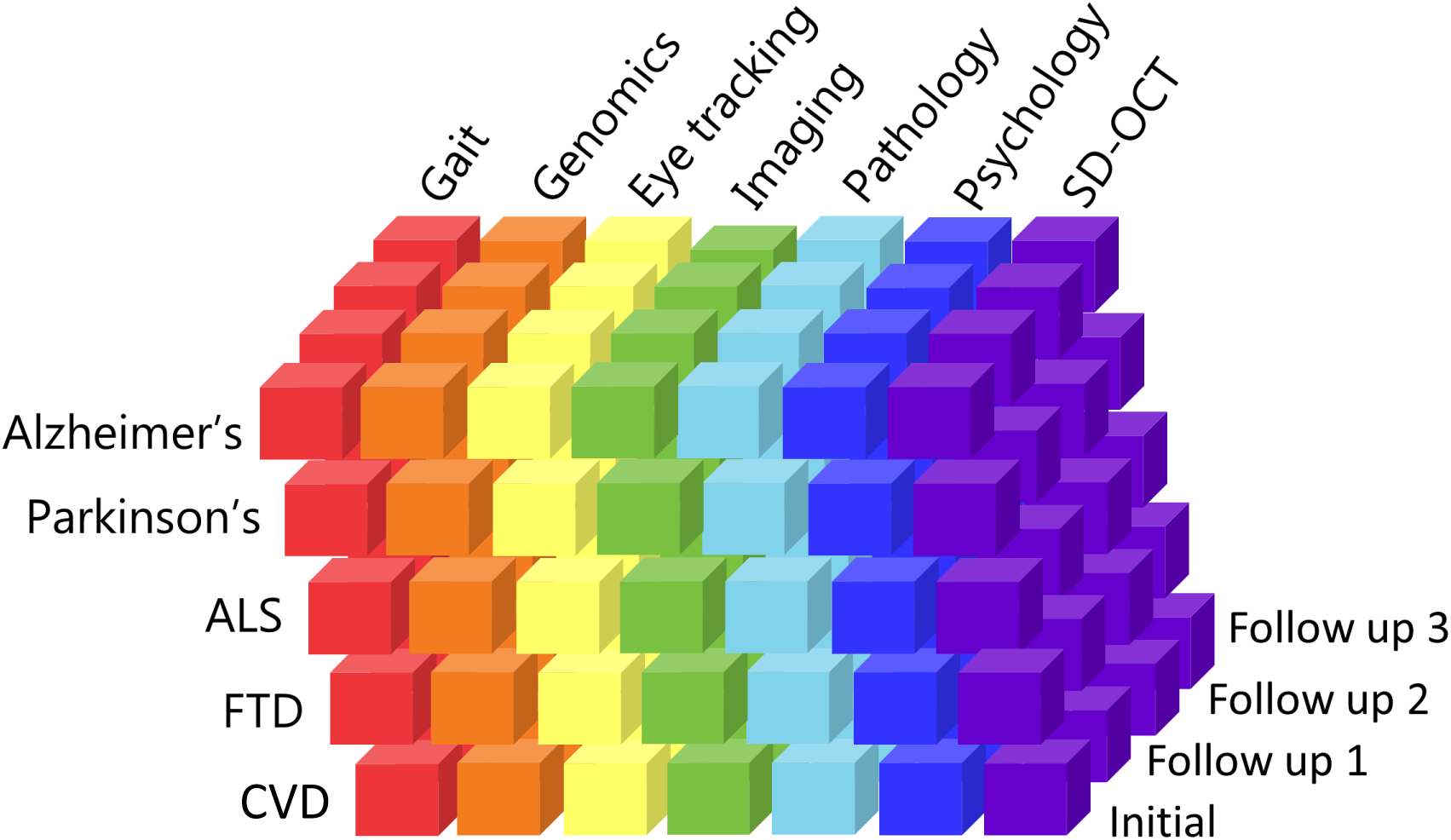
A visual representation of the data being collected in ONDRI, highlighting the inter-related and multi-dimensional nature of the disease groups, assessment platforms and time points.

### 1.1 Challenges Associated with Large Multi-site Imaging Studies

Multi-site imaging studies benefit from the standardization of imaging protocols to reduce the variance associated with imaging measurements. Efforts at protocol standardization have been undertaken or proposed, not only for AD and MCI as in ADNI (5), but also in psychosis in youth (8), multiple sclerosis (9), brain tumors (10), stroke (11), and traumatic brain injury (12) and depression (13). Several groups have also analyzed the measurable differences associated with scanning at various sites and scanners examining the effects of how vendor, field strength, sequence, and coil have the potential to impact the data. Remarkably there is good evidence to suggest that despite this variability, the data are sufficiently robust to provide comparable results across all of these conditions (14–18).

Obtaining images of sufficient subjective quality, free of motion artifacts, covering full fields of view, and free of other scanner-related artifacts are all critical in ensuring the future usability and ability to process the data through various pipelines. Certain sequences are more susceptible to motion and other artifacts (such as resting-state fMRI and DTI). Other sequences are of such paramount importance (e.g. T_1_-weighted imaging) that tolerance for motion is low as it will severely impact the quality of the data, and the usability of other sequences which are typically co-registered to T_1_-weighted anatomical images. Common practice to identify such artifacts or other issues is to use a manual visual review of all images to determine if their quality is sufficient. This requires expertly trained reviewers and guidelines, and although laborious, is a critical step in ensuring proper data quality. The introduction of images containing these artifacts could seriously impact the overall integrity of the results obtained, and make it difficult to detect more subtle but potentially important artifacts using automated quality control procedures. Additionally, timely review affords the opportunity in some cases to recall the particpant for repeat imaging, if the acquisition window has not been exceeded. A few examples of such manual quality control guidelines exist online, including a helpful manual from the group at the Harvard Centre for Brain Science (http://cbs.fas.harvard.edu/usr/mcmains/CBS_MRI_Qualitative_Quality_Control_Manual.pdf). For comparison, all ADNI scans were reviewed by a quality control team at the Mayo clinic for issues such as protocol deviation or motion artifact (19), similar to what is being done within ON-DRI.

### 1.2 Neuroinformatics

The processes and infrastructure involved in data capture, assessment, archive and databasing is often collectively referred to as informatics. Informatics relating to aspects of the brain is commonly referred to as neuroinformatics (20,21). Many tools and infrastructures are available for the purposes of data management and analytics, examples of which include NeuroGems (http://www.neurogems.org/), LORIS (Longitudinal Online Research and Imaging System) (22), XNAT (eXtenisble Neuroimaging Archive Toolkit) (23), fBIRN (Function Bioinformatics Research Network)(24), and COINS (25), or entire suites of software and hosting abilities such as NITRC (26) that provide a complete framework. Toolkits such as XNAT can also act as the image repository backbone for other interfaces such as SPReD (Stroke Patient Recovery Research Data-base-developed by the Heart and Stroke Foundation Canadian Partnership for Stroke Recovery; http://www.canadianstroke.ca/) (27), PURE-MIND (28), and Brain-CODE for ONDRI. Other data capture and informatics platforms such as Open Clinica (https://www.openclinica.com/), REDCap (http://projectredcap.org/) and RedMine (http://www.redmine.org/) are highly beneficial for the capture and management of non-imaging data such as neuropsychological assessments, other case-report forms, and managing daily operations.

There is currently still no consensus on the steps that should be undertaken to ensure high quality multi-site imaging data acquisition and management. Therefore, the purpose of the current manuscript is to share the processes and experiences of the ONDRI project to benefit the broader imaging and neuroinformatics communities.

## 2. METHODOLOGY

The ONDRI neuroinformatics framework consists of a number of software tools, pipelines and procedures designed to ensure high quality data acquisition, databasing, archiving, assessment, analysis, and tracking, an overview of which is shown in Figure 2. The primary platform for this set of tools is SPReD/XNAT through Brain-CODE. In addition to the MR imaging data being captured and managed through SPReD/XNAT (as well as data acquired through the eye tracking, ocular imaging and gait assessment platforms), other study-related data are captured with RedCap and tracked with RedMine, however, discussion of those will not be included here as they are beyond the scope of the current paper which solely describes the neuroimaging tools. Additionally, a commercial visualization ‘dashboard’ called Spotfire (http://spotfire.tibco.com/) is used to publish aggregated data tracking and analytics results to the web (see Figure 3). The pipelines and procedures that have been implemented in ONDRI are now described.

**Figure 2:**
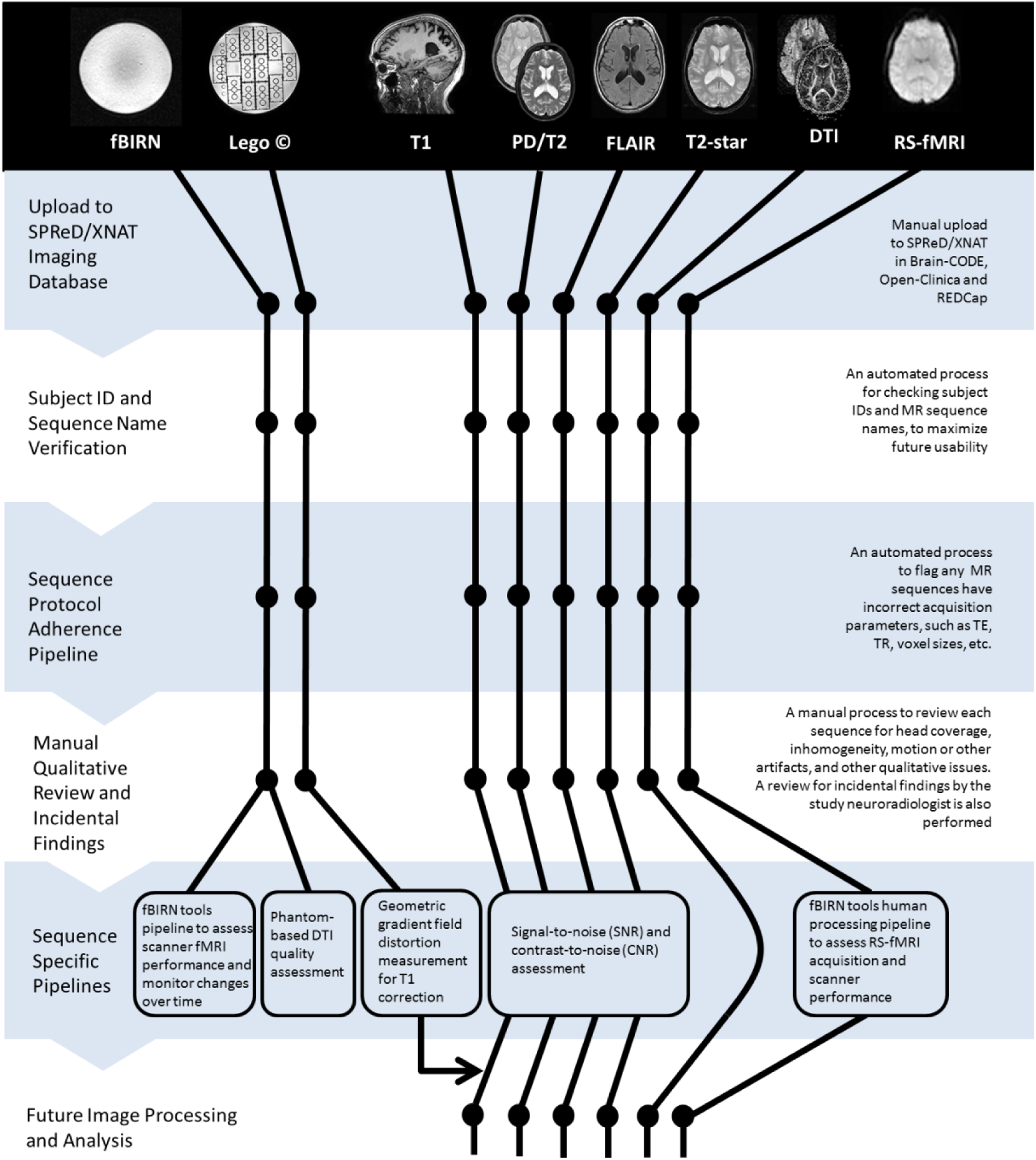
An overview of the pipelines and procedures for assessing MR imaging in ONDRI.

**Figure 3:**
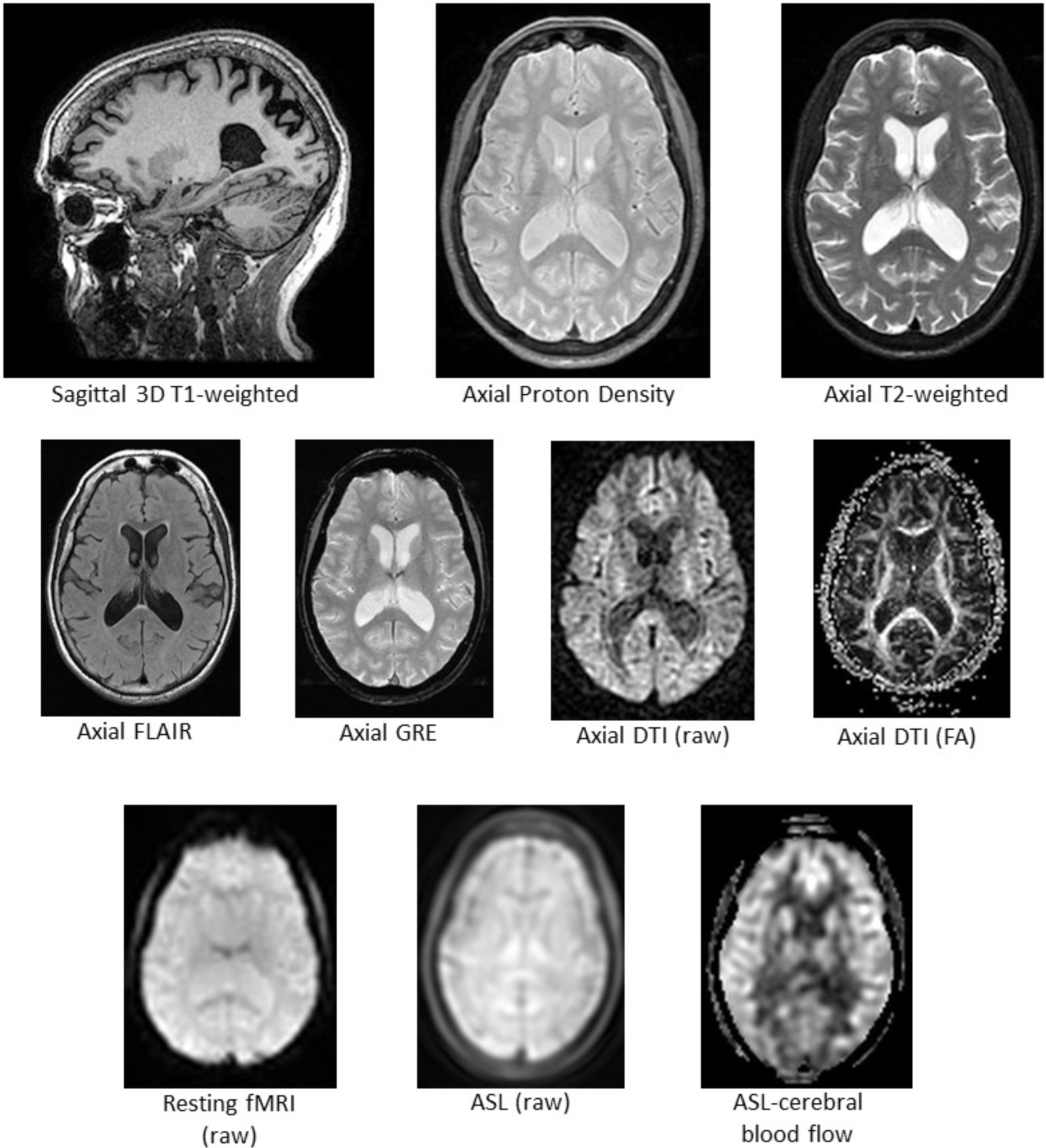
Examples of the MR imaging acquired for each participant (ASL at 1 site only)

### 2.1 Establishing the MRI Acquisition Parameters Across Sites

One of the primary goals of the ONDRI neuroimaging platform was to ensure consistent and compatible MRI data acquired between sites with different scanners using a standardized MRI protocol that would be adopted by several studies taking place across Canada, termed the Canadian Dementia Imaging Protocol (29,30). Collaboratively developed by a group of physicists, physicians and researchers from across Canada, the protocol has also been implemented in a multitude of projects including the Canadian Alliance for Healthy Hearts and Minds (CAHHM - http://cahhm.mcmaster.ca/); the Consortium d’Identification de la Maladie d’Alzheimer – Québec (CIMAQ - http://www.cima-q.ca/); the O2 study from the Consortium Québécois de Découverte du Médicament (http://www.cqdm.org/en/); the Medical Imaging Trials Network of Canada (MITNEC - http://www.mitnec.org/), and the Canadian Consortium for Neurodegeneration and Aging (CCNA; https://ccna-ccnv.ca/). The protocol is inspired by the standardized imaging protocols in ADNI-2, with some additional sequences added, and has been further harmonized across the three main MR vendor platforms including General Electric (GE), Philips and Siemens, to deliver comparable data from each scanner for both basic structural and advanced imaging sequences.

The CDIP protocol includes a series of advanced imaging sequences with high utility across many disease states including but not limited to stroke, dementia, neurodegeneration, traumatic brain injury (TBI), amyotrophic lateral sclerosis (ALS) and Parkinson’s disease. The protocol includes a high-resolution 3D isotropic T_1_-weighted scan for assessing fine anatomical detail and cortical thickness mapping (3DT1), an interleaved proton-density/T_2_ weighted image (PD/T2) for reliable skull-stripping and lesion assessment in deep cerebral nuclei and peri-vascular spaces, a fluid-attenuated inversion recovery (FLAIR) image for quantification and assessment of small vessel disease (SVD, e.g. white matter hyperintensities), a T_2_-star weighted gradient-echo sequence (T2GRE) for identification of microbleeds, 30 direction diffusion tensor imaging (DTI) for assessment of microstructural and white matter integrity, and resting state Blood Oxygen Level Dependent (BOLD) functional MRI for assessment of brain network connectivity (RS-fMRI) (Figure 4).

**Figure 4:**
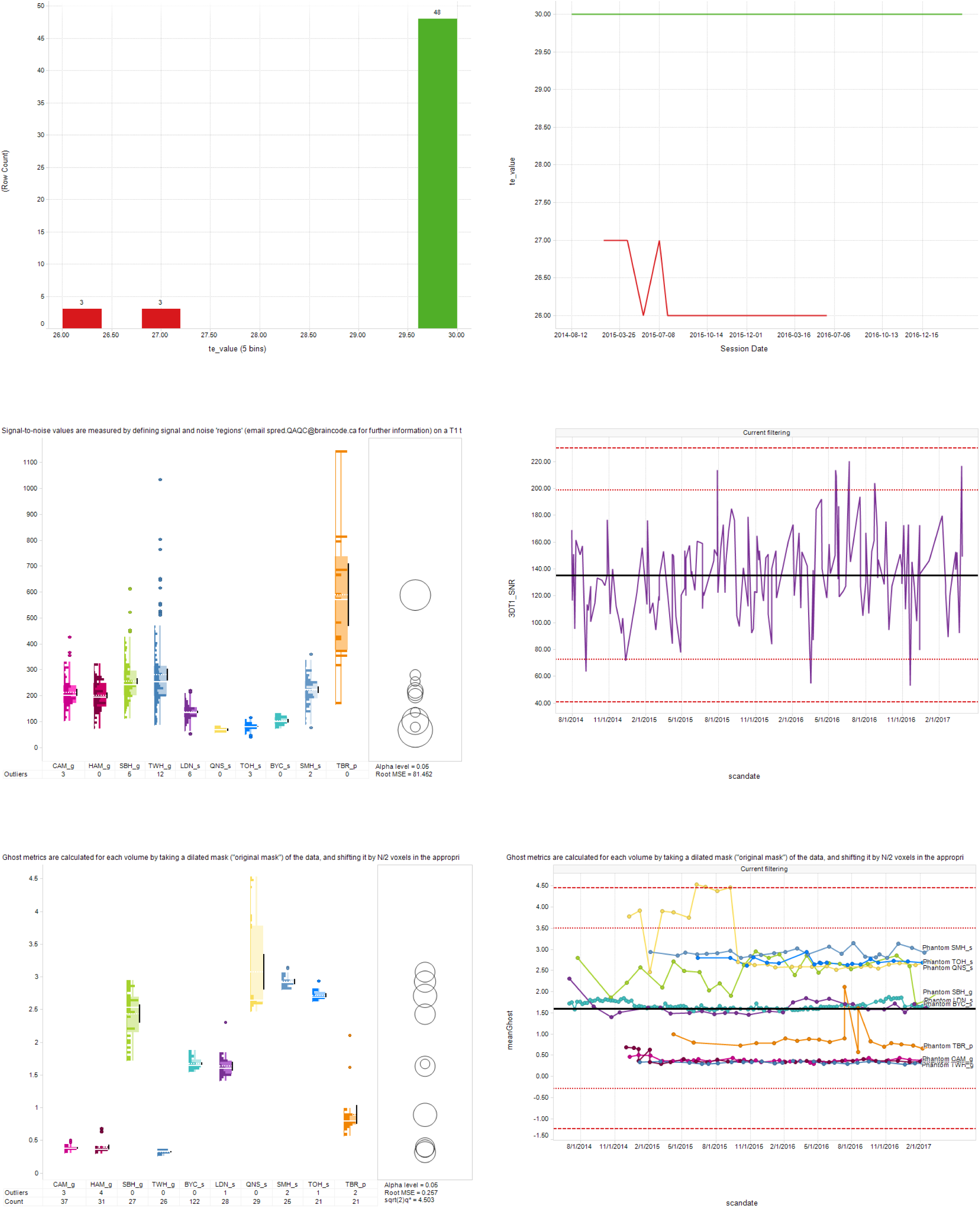
Sample images from the MR Quality Assurance (QA) and Control (QC) Spotfire dashboards. **Top panel**: Cross-sectional (left) and longitudinal (right) displays of the Scan Acquisition Parameter QA Dashboard depicting the Resting State fMRI TE values acquired from 54 scans from one ONDRI site. Green and red traces represent scans with TE values that were within and outside of the expected range, respectively. **Middle panel**: Cross-sectional display (left) of the Structural QC Pipeline Dashboard depicting signal-to-noise ratio (SNR) results of 3DT1 scans obtained at each of the ONDRI sites. Right panel depicts the longitudinal display of the same results for one site. Dashed and dotted horizontal red lines indicating ±3 and 2 standard deviation limits, respectively, of the site’s overall SNR average. **Bottom panel**: Cross-sectional (left) display of the fBIRN Phantom QA Results Dashboard depicting the ‘Ghosting value’ results from the monthly Resting State fMRI scans of the fBIRN phantom. Right panel depicts the longitudinal display of those results from each of the ONDRI scanners (separate lines), with dashed and dotted horizontal red lines indicating ±3 and 2 standard deviation limits, respectively, of the overall datapoint values.

The ONDRI protocol (Supplementary Table 1) was provided to the MRI physicists identified at each of the ten 3 Tesla MRI centres across the province of Ontario and included the following systems: a General Electric (GE, Milwaukee, WI) Discovery 750 was used at Sunnybrook Health Sciences Centre, McMaster University/Hamilton General Hospital, and the Centre for Addiction and Mental Health; a GE Signa HDxt at Toronto Western Hospital; a Philips Medical Systems (Philips, Best, Netherlands) Achieva system at Thunder Bay Regional Health Sciences Centre; Siemens Health Care (Siemens, Erlangen, Germany) Prisma at Sunnybrook Health Sciences Centre and London Health Sciences Centre/Parkwood Hospital; a Siemens TrioTim at Ottawa Hospital/Élisabeth Bruyère Hospital, Hotel Dieu Hospital/ Providence Care Hospital and Baycrest Health Sciences; and a Siemens Skyra at St. Michael’s Hospital.

**Table 1.** —part 1—ONDRI scan acquisition parameter ranges according to site

| Scan Type<br>Parameter value | Parameter ranges used in Acquisition Checker according to site and scanner type |  |  |  |  |
| --- | --- | --- | --- | --- | --- |
|  | CAM | MCM | SBH | TWH | BYC |
|  | (GE 3.0 Tesla Discovery MR750) | (GE 3.0 Tesla Discovery MR750) | (GE 3.0 Tesla Discovery MR750) | (GE 3.0 Tesla Signa HDxt) | (Siemens 3.0 Tesla Trio Tim) |
| 3DT1 |  |  |  |  |  |
| TR | [6.652 : 6.652] | [8.156 : 8.156] | [8.156 : 8.156] | [6.9 : 7.3] | [2300 : 2300] |
| TE | [2.928 : 2.928] | [3.18 : 3.18] | [3.18 : 3.18] | [2.8 : 3.1] | [2.98 : 2.98] |
| T1 | [400 : 400] | [400 : 400] | [400 : 400] | [400 : 400] | [900 : 900] |
| Flip | [11 : 11] | [11 : 11] | [11 : 11] | [11 : 11] | [9 : 9] |
| pixelBandwidth | [244.141 : 244.141] | [244.141 : 244.141] | [244.141 : 244.141] | [244.141 : 244.141] | [238 : 238] |
| Matrix Size | [256x256 : 256x256] | [256x256 : 256x256] | [256x256 : 256x256] | [256x256 : 256x256] | [256x256 : 256x256] |
| Voxel Size | [1x1x1 : 1x1x1] | [1x1x1 : 1x1x1] | [1x1x1 : 1x1x1] | [1x1x1 : 1x1x1] | [1x1x1 : 1x1x1] |
| slice | [176 : 176] | [176 : 176] | [176 : 176] | [176 : 176] | [176 : 176] |
| PD/T2 |  |  |  |  |  |
| TR | [3000 : 3000] | [3000 : 3000] | [3000 : 3000] | [3000 : 3000] | [3000 : 3000] |
| MultiEcho_TE1 | [10.2 : 11.2] | [10.3 : 11.1] | [10.7 : 12.0] | [10.3 : 10.7] | [10.5 : 11.5] |
| MultiEcho_TE2 | [83 : 89] | [83 : 89] | [88 : 95] | [84 : 86] | [86 : 90] |
| Flip | [125 : 125] | [125 : 125] | [125 : 125] | [90 : 90] | [165 : 165] |
| pixelBandwidth | [162.734 : 162.734] | [162.734 : 162.734] | [150.234 : 150.234] | [162.734 : 162.734] | [181 : 181] |
| Matrix Size | [256x256 : 256x256] | [256x256 : 256x256] | [256x256 : 256x256] | [256x256 : 256x256] | [256x256 : 256x256] |
| Voxel Size | [0.9375x0.9375x3 : 0.9375x0.9375x3] | [0.9375x0.9375x3 : 0.9375x0.9375x3] | [0.9375x0.9375x3 : 0.9375x0.9375x3] | [0.9375x0.9375x3 : 0.9375x0.9375x3] | [0.9375x0.9375x3 : 0.9375x0.9375x3] |
| slice | [96 : 96] | [96 : 96] | [96 : 96] | [96 : 96] | [96 : 96] |
| 2D FLAIR |  |  |  |  |  |
| TR | [9000 : 9000] | [9000 : 9000] | [9000 : 9000] | [9002 : 9002] | [9000 : 9000] |
| TE | [139 : 144] | [138 : 146] | [140 : 152] | [137 : 143] | [118 : 124] |
| T1 | [2250 : 2250] | [2250 : 2250] | [2250 : 2250] | [2250 : 2250] | [2500 : 2500] |
| Flip | [125 : 125] | [125 : 125] | [125 : 125] | [90 : 90] | [165 : 165] |
| pixelBandwidth | [195.312 : 195.312] | [195.312 : 195.312] | [217.031 : 217.031] | [195.312 : 195.312] | [222 : 224] |
| Matrix Size | [256x256 : 256x256] | [256x256 : 256x256] | [256x256 : 256x256] | [256x256 : 256x256] | [192x256 : 192x256] |
| Voxel Size | [0.9375x0.9375x3 : 0.9375x0.9375x3] | [0.9375x0.9375x3 : 0.9375x0.9375x3] | [0.9375x0.9375x3 : 0.9375x0.9375x3] | [0.9375x0.9375x3 : 0.9375x0.9375x3] | [0.9375x0.9375x3 : 0.9375x0.9375x3] |
| slice | [48 : 48] | [48 : 48] | [48 : 48] | [48 : 48] | [48 : 48] |
| T2-star* | *_PMRI | *_PMRI | *_PMRI |  |  |
| TR | [650 : 650] | [650 : 650] | [650 : 650] | [650 : 650] | [650 : 650] |
| TE | [20 : 20] | [20 : 20] | [20 : 20] | [20 : 20] | [20 : 20] |
| Flip | [20 : 20] | [20 : 20] | [20 : 20] | [20 : 20] | [20 : 20] |
| pixelBandwidth | [150.234 : 150.234] | [150.234 : 150.234] | [150.234 : 150.234] | [150.234 : 150.234] | [199 : 199] |
| Matrix Size | [256x256 : 256x256] | [256x256 : 256x256] | [256x256 : 256x256] | [256x256 : 256x256] | [256x256 : 256x256] |
| Voxel Size | [0.9375x0.9375x3 : 0.9375x0.9375x3] | [0.9375x0.9375x3 : 0.9375x0.9375x3] | [0.9375x0.9375x3 : 0.9375x0.9375x3] | [0.9375x0.9375x3 : 0.9375x0.9375x3] | [0.9375x0.9375x3 : 0.9375x0.9375x3] |
| slice | [192 : 192] | [188 : 196] | [188 : 196] | [48 : 48] | [48 : 48] |
| fMRI-RS |  |  |  |  |  |
| TR | [2400 : 2400] | [2400 : 2400] | [2400 : 2400] | [2400 : 2400] | [2400 : 2400] |
| TE | [30 : 30] | [30 : 30] | [30 : 30] | [30 : 30] | [30 : 30] |
| Flip | [70 : 70] | [70 : 70] | [70 : 70] | [70 : 70] | [70 : 70] |
| pixelBandwidth | [7812.5 : 7812.5] | [7812.5 : 7812.5] | [7812.5 : 7812.5] | [7812.5 : 7812.5] | [2441 : 2441] |
| Matrix Size | [64x64 : 64x64] | [64x64 : 64x64] | [64x64 : 64x64] | [64x64 : 64x64] | [448 : 448] |
| Voxel Size | [3.5x3.5x3.5 : 3.5x3.5x3.5] | [3.5x3.5x3.5 : 3.5x3.5x3.5] | [3.5x3.5x3.5 : 3.5x3.5x3.5] | [3.5x3.5x3.5 : 3.5x3.5x3.5] | [3.5x3.5x3.5 : 3.5x3.5x3.5] |
| slice | [10250 : 10250] | [10250 : 10250] | [10000 : 10000] | [10250 : 10250] | [250 : 250] |
| DTI |  |  |  |  |  |
| TR | [9000 : 9000] | [9000 : 9000] | [9000 : 9000] | [11700 : 11700] | [9400 : 9400] |
| TE | [82 : 89] | [81 : 90] | [80 : 95] | [105 : 110] | [94 : 99] |
| Flip | [90 : 90] | [90 : 90] | [90 : 90] | [90 : 90] | [90 : 90] |
| pixelBandwidth | [3906.25 : 3906.25] | [3906.25 : 3906.25] | [3906.25 : 3906.25] | [3906.25 : 3906.25] | [2056.0 : 2056.0] |
| Matrix Size | [128x128 : 128x128] | [128x128 : 128x128] | [128x128 : 128x128] | [128x128 : 128x128] | [1152x1152 : 1152x1152] |
| Voxel Size | [2x2x2 : 2x2x2] | [2x2x2 : 2x2x2] | [2x2x2 : 2x2x2] | [2x2x2 : 2x2x2] | [2x2x2 : 2x2x2] |
| slice | [2310 : 2310] | [2310 : 2310] | [2310 : 2310] | [2310 : 2310] | [31 : 31] |
| DTI_b0_1 |  |  |  |  |  |
| TR | - | - | - | - | [9400 : 9400] |
| TE | - | - | - | - | [94 : 99] |
| Flip | - | - | - | - | [90 : 90] |
| pixelBandwidth | - | - | - | - | [2056.0 : 2056.0] |
| Matrix Size | - | - | - | - | [128x128 : 128x128] |
| Voxel Size | - | - | - | - | [2x2x2 : 2x2x2] |
| slice | - | - | - | - | [70 : 70] |
| DTI_b0_2 |  |  |  |  |  |
| TR | - | - | - | - | [9400 : 9400] |
| TE | - | - | - | - | [94 : 99] |
| Flip | - | - | - | - | [90 : 90] |
| pixelBandwidth | - | - | - | - | [2056.0 : 2056.0] |
| Matrix Size | - | - | - | - | [128x128 : 128x128] |

**Table 1.**
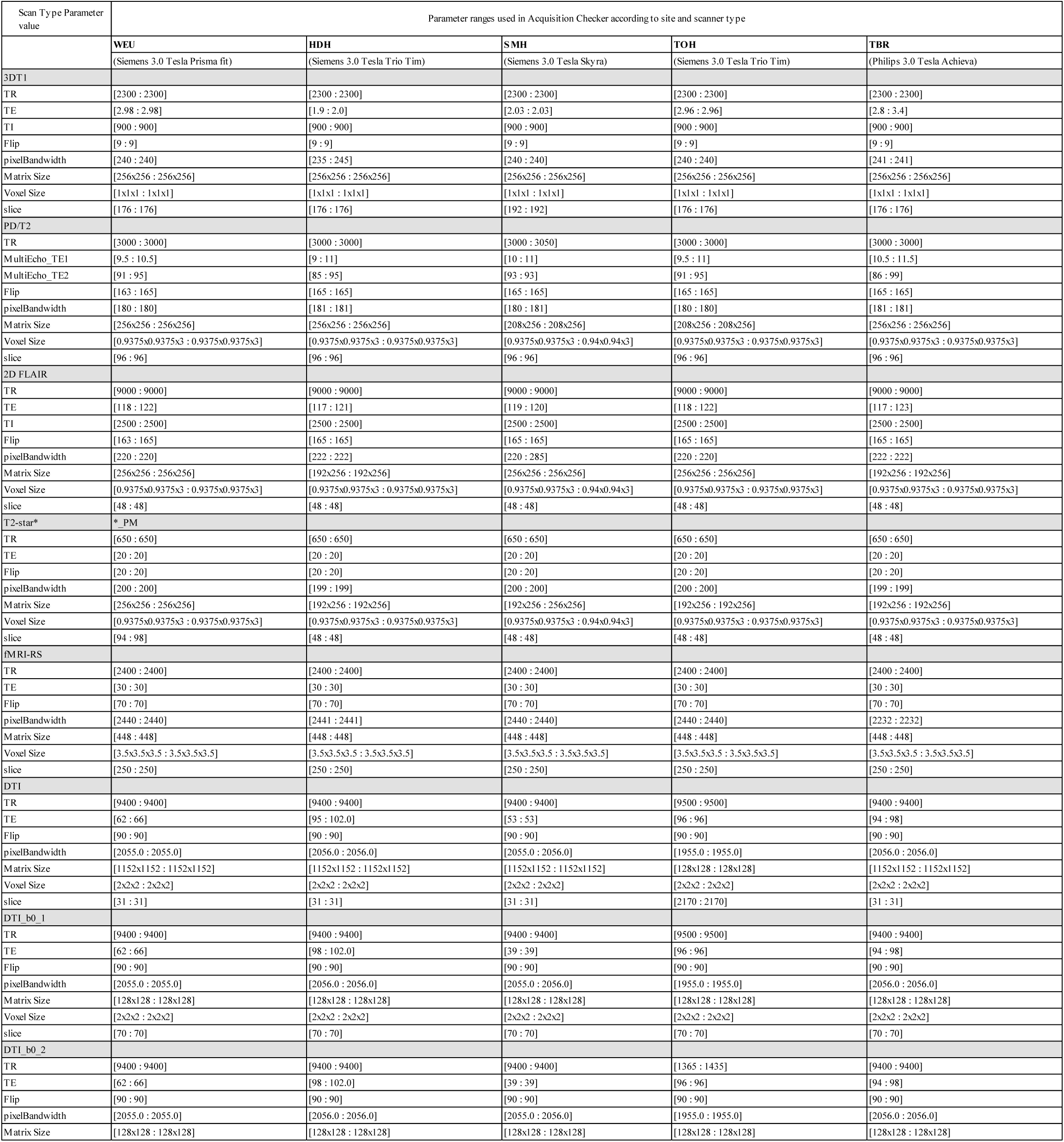
—part 2—ONDRI scan acquisition parameter ranges according to site

Each site acquired a series of test images that were sent to the ONDRI imaging platform for verification. Based on instructions from the imaging platform, adjustments were made iteratively at each site until all sequences achieved similar qualitative image quality and adhered to the ONDRI protocol after which the site was allowed to scan participants recruited to the ONDRI study.

Below, quality assurance (QA) refers to processes performed to ensure the procedures, processes and systems put in place to collect data were functioning optimally, whereas quality control (QC) refers to steps taken applied to directly ascertain the quality of the collected participant data.

### 2.2 Automatic naming convention adherence pipeline

Within an OBI research program (“Integrated Discovery Program” or IDP), each participant has a uniquely assigned Participant Identification Code, comprised of a three-letter Program Code, two-digit Study Number, three-letter Site Identification Code (usually corresponding to recruitment site), and four-digit participant number assigned by ONDRI clinical managers (e.g., ‘OND01_SBH_0001’). Data associated with a given experimental session are then uploaded into the appropriate site-specific project for ethics-related access control in a participant directory in SPReD/XNAT as independent files within a uniquely named session folder with a participant ID prefix, followed by a two-digit visit ID, and a session ID denoted by the letters “SE” and a two-digit session number, followed by a modality code (e.g., ‘OND01_SBH_0001_01_SE01_N’ for neuroimaging).

Proper adherence to this naming convention is essential in order for Brain-CODE to associate imaging data with those collected from other platforms (e.g., demographic and clinical measures, genomics, etc.). As well, adherence to the naming convention also enables ONDRI data to be federated with data collected from any other Brain-CODE IDP. The naming consistency QC pipeline is a Python executable that iterates nightly through the data uploaded to SPReD/XNAT in Brain-CODE and checks whether the names of the uploaded files comply with the naming convention. If non-compliance is detected, within 24 hours the data uploader is notified, and after 7 days the IDP Program Manager and a member of the SPReD Administration staff are also notified by daily email until the naming problem is corrected.

In addition to the automated pipelines above, manual checks of data records are also undertaken periodically. For example, dates of data acquisition as recorded in SPReD are checked against those recorded in the REDCap clinical assessment records in order to flag discrepancies and catch potential entry errors. Because most metadata associated with a given neuroimaging session are extracted directly from the DICOM fields themselves, such errors are relatively low relative to the other (non-DICOM) SPReD sessions that rely on manual metadata entry (e.g., eye tracking, ocular imaging, gait and balance assessments). Nevertheless, date errors can and inevitably do arise and therefore need to be checked in order to avoid possible complications downstream (e.g., a dataset being excluded from analysis because it was deemed to be acquired outside of the allotted time window).

### 2.3 Automatic MR protocol parameter adherence pipeline

Ensuring that a particular imaging protocol is strictly adhered to at any given acquisition site is critical when intending to pool data (31). Despite best efforts to create harmonized imaging protocols that will ideally remain unchanged, various factors can introduce minor or major deviations to the protocol. These factors can include technologist/operator decision making and inputs, software or hardware upgrades that can reset various parameters, or even the scanners themselves making automatic on-the-fly adjustments (e.g., to account for SAR - Specific Absorption Rate) to adjust for patient weight or ambient temperature in the scanner room. Dozens of specific parameters, which are fundamental to each MR sequence, are encoded into the metadata DICOM header tags. These can be extracted, checked against a site/vendor/model specific protocol template and flagged if variations are observed. Being able to quickly and automatically identify and track these parameters is essential in identifying occasional or ongoing protocol deviations and determining if it will have a detrimental effect on the quality of the acquired data. Such checking can be accomplished through the use of an automatic protocol adherence pipeline.

The Brain-CODE Scan Protocol QC pipeline is a Python executable that runs nightly and compares the parameters for all newly uploaded scans within an MRI session on SPReD against a reference protocol defined by the relevant IDP. The protocols are configured on a project-by-project and scanner-by-scanner basis for each scanning site. Each protocol file defines a set of pulse sequences that are to be acquired for every session, along with a set of values for the acquisition parameters for each sequence. Each parameter value has an upper and lower value against which the actual scan parameters are evaluated, and historical results are aggregated and displayed using Spotfire (upper panel of Figure 3a). Parameter failures or missing/ incorrectly named scans are emailed out to the technologists at each site as well as the central QC team. Within 24 hours of a parameter failure occurring and being logged, the central QC team determines whether the deviation represents an unacceptable variation and if necessary, contacts and works with the scanning site to try to ascertain and correct the cause of the failure.

### 2.4 Automatic signal-to-noise (SNR) and contrast-to-noise (CNR) quantitative pipelines

Pipelines have been implemented to produce statistical measures to determine the quality of the uploaded ONDRI structural MR images. The Structural QC pipeline automatically registers every newly uploaded high-resolution T1 scan to the MNI152 template. This multi-stage registration is performed using the ANTs (Advanced Normalization Tools) package. Following registration, the signal-to-noise (SNR) of the left and right putamen as defined by the LPBA40 atlas (http://www.loni.usc.edu/atlases/Atlas_Detail.php?atlas_id=12) is automatically measured. Signal is measured as the mean intensity value of the LPBA40 putamen regions, and noise by the standard deviation of noise spheres well outside the brain in the MNI152 space, to the anterior left and anterior right of the skull. Similarly, contrast-to-noise (CNR) is measured as the difference between the mean left putamen signal and the mean right putamen signal, divided by the standard deviation of the noise spheres. Four structural MR scans (PD/T2, 3DT1, T2-star (GRE), and 2D FLAIR) are processed to determine the SNR and CNR values, which are used as QC measures and indicators of scan acquisition quality at each ONDRI site.

Results from this pipeline are summarized in a QC Assessment file stored with each ONDRI MR session on SPReD. These results are also aggregated regularly and displayed on a Spotfire dashboard (see middle panel of Figure 3). The visualization of QC data on this and subsequent dashboards is used by the QA/QC team to monitor site performance over time and flag parameter values that exceed critical threshold values >3 standard deviations (SD) from the scanner’s historical mean (n.b., ‘anomalous’ datapoints > 4SD are excluded from the historical mean calculation in order to avoid artificially inflating the threshold limits).

### 2.5 FreeSurfer pipeline

The FreeSurfer pipeline is a Python-based script that iterates nightly through the ONDRI imaging repository for newly uploaded T1 scans. When found, the pipeline executes the “recon-all” command from the FreeSurfer Software Suite (version 6.0.0) to perform all FreeSurfer cortical reconstruction processes including subcortical segmentation, cortical parcellation, brainmask generation, as well as the generation ofng statistical outputs from each step, including Euler number. (For a full description of the FreeSurfer pipeline, see https://surfer.nmr.mgh.harvard.edu/fswiki/recon-all. The resulting output files are written to the original T1 scan folder in the SPReD/XNAT imaging data repository where they are made available to researchers for subsequent data processing and quality checks.

### 2.6 BIDS conversion

An important mandate of the OBI and its IDP-funded research is a commitment to an “Open Data Interface” in which data are to be made available to the global research community in a manner that is in keeping with the FAIR Data Principles of Findable, Accessible, Interoperable and Reusable (32,33). In recent years a concerted effort to develop a simple standardized method of organizing, annotating and describing neuroimaging data has resulted in the emergence of the Brain Imaging Data Structure (BIDS) (34). Currently adopted by databases such as OpenNeuro, SchizConnect, Developing Human Connectome Project, and FCP-INDI, BIDS is largely based on a formalized file/folder structure with controlled vocabulary and JSON-based metadata files.

The BIDS conversion pipeline (XNAT2BIDS converter) is a script that automatically converts a python script built on a modified version of Nipy’s heudiconv package (https://github.com/nipy/heudiconv) which operates by retrieving data from SPReD’s XNAT MR sessions and converting those data to the Brain Imaging Data Structure (BIDS). Manual QC reports and other meta information associated with each session are stored as sidecar JSON files. A validation script is then executed on the resulting BIDS folder to ensure data integrity and that no missing values exist. After a final validation step, the ‘BIDSified’ data are uploaded back into the original SPReD session, under the Resource Files.

### 2.7 Pipeline for monitoring and correction of MR scanner geometric gradient field distortions with the Lego® phantom

Geometric distortions in anatomical MR images caused by gradient magnetic field non-linearity are a major source of measurement variability in morphometric analyses of human brain structures. This increased variability reduces the statistical power to detect changes in cross-sectional and longitudinal studies. Every MRI centre in ONDRI estimates and corrects these geometric distortions using data acquired at regular intervals from a LEGO® phantom (35,36) as a part of QA procedures. The coefficients of the coordinate mapping functions, which are defined using spherical harmonic expansion, are calculated using the apparent displacements of corresponding points relative to the known ground truth of the phantom (35). The estimated coordinate mapping function for each ONDRI imaging centre is then used to correct geometric distortions in the 3DT1 MR images which are made available for future analyses.

### 2.8 Automatic DTI quality assessment pipeline

DTI is sensitive to artifacts and distortions which degrade image quality, compromising the accuracy and sensitivity of outcome metrics. DTI image quality is inherently degraded by three main effects: (i) B_0_ field inhomogeneity, (ii) eddy currents, and (iii) Nyquist ghosting. Both acquisition-based and post-processing approaches can compensate for these effects to varying degrees, depending primarily on the specific scanner, vendor and method. This can lead to a large variation in data quality across sites, causing site-specific biases. Thus, there is a need for a phantom based DTI-QA tool. We have used such a tool applied to the homogeneous, agar-filled spherical phantom (i.e., fBIRN phantom (17,24)) scanned in the ONDRI study, to characterize and track systematic differences in DTI metrics across sites.

The DTI-QA tool employed in this study (37) uses the fact that all aforementioned effects occur in the phase encode (PE) rather than the frequency encode (FE) direction. The B_0_ field inhomogeneity, eddy current, and Nyquist ghosting effects are quantified by the following metrics respectively: (i) circular asymmetry of non-DWIs measured by the ratio of diameters in PE and FE directions, (ii) pixel shift differences in phantom outline (in PE direction) between each DWI (diffusion-weighted image) and the first non-DWI, and (iii) ratio of average signal in ROIs outside the phantom in PE and FE directions. In addition, signal-to-noise-ratio (SNR) of individual images is also calculated as a measure for image quality and scanner stability. Finally, fractional anisotropy (FA) maps are computed using FSL (FMRIB’s Software Library).

### 2.9 Automatic Resting State fMRI scanner performance monitoring pipelines (fBIRN)

To monitor the performance of the MR scanners, OBI-funded fBIRN phantoms (17cm diameter, agar-filled spheres from the Biomedical Informatics Research Network) were purchased for each site, and scanned by site members at approximately monthly intervals using the ONDRI resting state pulse sequence (17). Acquired data from each site was uploaded to SPReD within 24-48 hours of acquisition, and automatically processed within the following 24 hours using the fBIRN pipeline software (Biomedical Informatics Research Network: www.nbirn.net). The fBIRN pipeline creates a full QA report (i.e., index.html) that is stored in the session’s fBIRN Phantom Results folder on SPReD/XNAT, and is available for download (see sample QA report in Appendix 1). Each report includes summary measures for each session such as the mean signal, the mean signal-to-noise ratio, drift, ghosting, etc. See (17,18) for a description of the pipeline and QA measures; the detailed parameter descriptions are available here: https://ww.nitrc.org/frs/download.php/275/fBIRN_phantom_qaProcedures.pdf, as well as in Appendix 2). A separate, nightly, python-based pipeline then extracts the summary variables from each QA report, and aggregates them into a comma separated variable (csv) file used to populate a Spotfire® dashboard for tracking the QA results over time. Notification thresholds for sites in which a given fBIRN parameter value exceeds 3 standard deviations from that site’s historical mean are calculated and displayed on the Spotfire® dashboard (see bottom panel of Figure 3). When an outlier is found in phantom data, an email is sent to the relevant MR personnel at the site and an fBIRN rescan is requested. If that rescanned phantom continues to show the parameter deviation, an alert will be sent to the ONDRI MR QC team who will follow up with the site’s MR lead to ascertain the problem. In addition, the MR team will examine any recently acquired human data from that scanner to determine whether a potential scanner issue has impacted those data.

Because of the challenges in obtaining and tracking the phantom scans, monthly scan reminders are sent out to each site’s MR technician or MR designate if that site’s previous fBIRN scan was 21-28 days prior. Overdue scan notifications are emailed on a weekly basis to those people as well as their site’s MR Principal Investigator if no fBIRN phantom sessions have been uploaded to SPReD for 30 days since the date of that site’s most recent acquisition on SPReD.

### 2.10 Manual procedures for image assessment

Manual visual assessment of image quality is performed by trained expert raters using the SPReD interface within 48 hours of image upload on all MRI data. Each imaging sequence is reviewed independently for quality including full-brain coverage (on a two-point scale - complete or incomplete), and motion and other image artifacts on a three-point scale (none, mild or severe). Imaging that is found to have insufficient coverage, excessive motion or other imaging artifacts that may interfere with future processing and usability are marked as questionable or unusable depending on severity and if flagged as unusable, are made unavailable for subsequent analyses.

Patients assessed to have poor quality 3DT1 imaging are asked to return for a rescan when possible; otherwise the patient’s data are subsequently not used, as the T1 forms the basis of the overall determination of a session’s ‘pass’ or ‘fail’.

In addition to the timely manual visual assessment, a neuroradiologist reviews every T1, PD/T2, FLAIR and T2-star sequence for incidental findings, as well as provides a microbleed count and white matter lesion rating (ie. Fazekas ratings).

## 3. RESULTS

### 3.1 Automatic naming convention adherence pipeline

The naming convention pipeline has been one of the single most important QA pipelines to date, ensuring that all data uploads conform to the standardized naming convention. Not surprisingly, filename errors were initially quite high and became less frequent over time as uploaders became accustomed to the naming convention. It is worth noting that, even years into the study, naming errors continue to occur, further underscoring the importance of this basic QA tool.

### 3.2 Automatic MR protocol parameter adherence pipeline

The Scan Acquisition Parameter Checker has been used to assess the parameter values from 989 sessions (including 15 reacquisition sessions), comprised of 1026 3DT1 scans, 1014 PD/ T2, 1059 2D Flair, 1194 T2-star, 1009 Resting State fMRI, and 988 DTI scans. Where applicable, parameter values included TR, TE, Ti, flip angle, pixel bandwidth, matrix size, voxel size, and number of slices. Table 1 lists the parameter ranges according to each acquisition site and scanner type. Parameter values that fell within the specified range were assigned a ‘pass” rating by the Scan Acquisition Parameter Checker, while those falling outside were flagged as a ‘fail’ and the QA/QC team was alerted to assess whether the parameter deviation was severe enough to render the scan unusable and/or to follow-up with the site’s MR personnel in order to avoid future scan parameter fails. Table 2 summarizes the overall Pass and Fail rate for each scan type. While adherence to expected values for any given parameter tended to be quite high within each pulse sequence (average ‘pass’ score for the 6-8 parameters that were tracked for each of the sequences was as follows: 3DT1 = 99.5%, PD/T2 = 98.9%, 2D Flair = 98.3%, T2-star = 99.8%, Resting State fMRI = 99.3%, DTI = 98.0%), the proportion of scans that had ‘pass’ ratings for all parameters within a given scan sequence was naturally lower. In particular, only 96.8% of 3DT1 scans, 87.1% of PD/ T2 scans, 87.9 % of 2D Flair scans, 95.6% of T2-star scans, 93.4% of rs-fMRI scans, and 77.5% of DTI scans, were not associated with any parameter warnings/errors. Each scan acquisition failure was scrutinized by the QC team and incorporated into the overall Usability rating. As can be seen in Table 2, the vast majority of parameter violations were ultimately deemed to not significantly impact the overall usability of the scan (e.g., many DTI scans acquired more than the expected number of slices) and, provided that they passed the Manual QC visual inspection, were given an overall scan Usability rating of ‘usable’ or ‘questionable’, rather than ‘unusable’.

**Table 2:** Proportion of adherence to scan parameter acquisition protocol

| Scan type | Scan Parameter Acquisition Checker Outcome (pass/fail) and ultimate scan usability rating once Manual QC team review was incorporated | Number of sessions | Mean proportion of protocol adherence (i.e., within acceptable range of parameter value) across all sites according to each parameter |  |  |  |  |  |  |  |
| --- | --- | --- | --- | --- | --- | --- | --- | --- | --- | --- |
|  |  |  | Tr | Te | Ti | flip | Pixel bandwidth | matrix size | voxel size | slice |
| 3DT1 | **pass** | **993** | **0.991** | **0.985** | **1** | **1** | **1** | **0.999** | **1** | **0.983** |
|  | questionable | 350 | 0.35 | 0.348 | 0.354 | 0.356 | 0.356 | 0.356 | 0.356 | 0.349 |
|  | unusable | 55 | 0.057 | 0.055 | 0.058 | 0.058 | 0.058 | 0.058 | 0.058 | 0.057 |
|  | usable | 588 | 0.585 | 0.583 | 0.587 | 0.587 | 0.587 | 0.586 | 0.587 | 0.578 |
|  | **fail** | **33** | **0.009** | **0.015** | **0** | **0** | **0** | **0.001** |  | **0.017** |
|  | questionable | 15 | 0.006 | 0.008 | 0 | 0 | 0 | 0 | 0 | 0.007 |
|  | unusable | 4 | 0.001 | 0.003 | 0 | 0 | 0 | 0 | 0 | 0.001 |
|  | usable | 14 | 0.002 | 0.004 | 0 | 0 | 0 | 0.001 | 0 | 0.009 |
| PDT2 | **pass** | **883** | **0.98** | - | - | **0.968** | **1** | **0.998** | **1** | **0.964** |
|  | questionable | 250 | 0.279 | - | - | 0.279 | 0.292 | 0.291 | 0.292 | 0.283 |
|  | unusable | 22 | 0.03 | - | - | 0.03 | 0.032 | 0.032 | 0.032 | 0.024 |
|  | usable | 611 | 0.672 | - | - | 0.66 | 0.677 | 0.676 | 0.677 | 0.658 |
|  | **fail** | **131** | **0.02** | - | - | **0.032** | **0** | **0.002** | **0** | **0.036** |
|  | questionable | 46 | 0.013 | - | - | 0.013 | 0 | 0.001 | 0 | 0.009 |
|  | unusable | 10 | 0.002 | - | - | 0.002 | 0 | 0 | 0 | 0.008 |
|  | usable | 75 | 0.005 | - | - | 0.017 | 0 | 0.001 | 0 | 0.019 |
| 2DFlair | **pass** | **931** | **0.982** | **0.96** | **0.981** | **0.972** | **0.991** | **0.998** | **1** | **0.965** |
|  | questionable | 410 | 0.453 | 0.432 | 0.452 | 0.44 | 0.456 | 0.463 | 0.464 | 0.455 |
|  | unusable | 73 | 0.076 | 0.076 | 0.076 | 0.077 | 0.077 | 0.077 | 0.077 | 0.072 |
|  | usable | 448 | 0.453 | 0.452 | 0.453 | 0.454 | 0.457 | 0.458 | 0.459 | 0.438 |
|  | **fail** | **128** | **0.018** | **0.04** | **0.019** | **0.028** | **0.009** | **0.002** | **0** | **0.035** |
|  | questionable | 81 | 0.01 | 0.031 | 0.011 | 0.024 | 0.008 | 0.001 | 0 | 0.008 |
|  | unusable | 9 | 0.002 | 0.002 | 0.002 | 0 | 0 | 0 | 0 | 0.006 |
|  | usable | 38 | 0.006 | 0.007 | 0.006 | 0.005 | 0.002 | 0.001 | 0 | 0.021 |
| T2star | **pass** | **1142** | **0.996** | **0.995** | - | **1** | **1** | **0.998** | **1** | **0.962** |
|  | questionable | 270 | 0.237 | 0.259 | - | 0.239 | 0.239 | 0.239 | 0.239 | 0.228 |
|  | unusable | 29 | 0.029 | 0.024 | - | 0.03 | 0.03 | 0.03 | 0.03 | 0.026 |
|  | usable | 843 | 0.729 | 0.712 | - | 0.731 | 0.731 | 0.729 | 0.731 | 0.709 |
|  | **fail** | **52** | **0.004** | **0.005** | - | **0** | **0** | **0.002** | **0** | **0.038** |
|  | questionable | 15 | 0.002 | 0.001 | - | 0 | 0 | 0 | 0 | 0.011 |
|  | unusable | 7 | 0.001 |  | - | 0 | 0 | 0 | 0 | 0.004 |
|  | usable | 30 | 0.002 | 0.004 | - | 0 | 0 | 0.002 | 0 | 0.023 |
| RS fMRI | **pass** | **942** | **0.998** | **0.984** | - | **1** | **0.992** | **0.998** | **0.988** | **0.962** |
|  | questionable | 410 | 0.425 | 0.416 | - | 0.426 | 0.422 | 0.426 | 0.419 | 0.424 |
|  | unusable | 41 | 0.067 | 0.067 | - | 0.067 | 0.066 | 0.067 | 0.067 | 0.041 |
|  | usable | 491 | 0.505 | 0.501 | - | 0.506 | 0.503 | 0.504 | 0.501 | 0.498 |
|  | **fail** | **67** | **0.002** | **0.016** | - | **0** | **0.008** | **0.002** | **0.012** | **0.038** |
|  | questionable | 20 | 0.001 | 0.011 | - | 0 | 0.004 | 0 | 0.007 | 0.002 |
|  | unusable | 27 |  | 0.001 | - | 0 | 0.001 | 0 | 0 | 0.027 |
|  | usable | 20 | 0.001 | 0.004 | - | 0 | 0.003 | 0.002 | 0.005 | 0.009 |
| DTI | **pass** | **766** | **0.928** | **0.963** | - | **1** | **0.997** | **0.997** | **0.996** | **0.859** |
|  | questionable | 326 | 0.385 | 0.405 | - | 0.42 | 0.42 | 0.42 | 0.42 | 0.366 |
|  | unusable | 26 | 0.043 | 0.046 | - | 0.048 | 0.047 | 0.047 | 0.047 | 0.032 |
|  | usable | 414 | 0.501 | 0.512 | - | 0.532 | 0.53 | 0.53 | 0.529 | 0.461 |
|  | **fail** | **222** | **0.072** | **0.037** | - | **0** | **0.003** | **0.003** | **0.004** | **0.141** |
|  | questionable | 89 | 0.035 | 0.015 | - | 0 | 0 | 0 | 0 | 0.054 |
|  | unusable | 21 | 0.005 | 0.002 | - | 0 | 0.001 | 0.001 | 0.001 | 0.015 |
|  | usable | 112 | 0.031 | 0.02 | - | 0 | 0.002 | 0.002 | 0.003 | 0.072 |

### 3.3 Automatic signal-to-noise (SNR) and contrast-to-noise (CNR) quantitative pipelines

As of the ‘new enrollment’ cut-off date, the Structural QC pipeline had been used to analyze 3904 structural scans from the ONDRI scanners. Table 3 summarizes the within-site normalized SNR and CNR values for each of the four structural MR scans (PD/T2, 3DT1, T2-star, and 2D FLAIR) of the analyzed data.

**Table 3.**
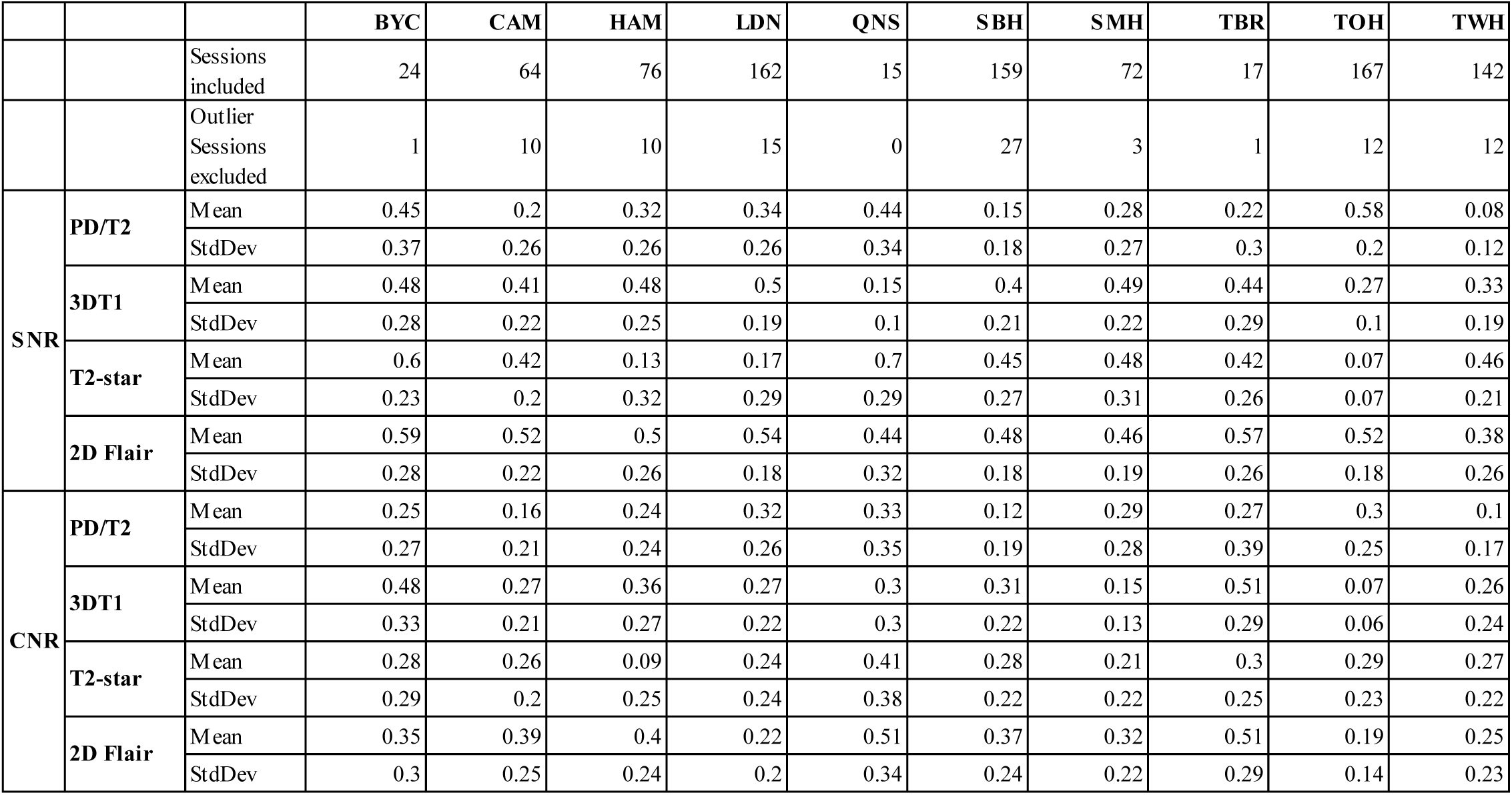
Structural QC Signal-to-noise (SNR) and contrast-to-noise (CNR) Results

|  |  |  | BYC | CAM | HAM | LDN | QNS | SBH | SMH | TBR | TOH | TWH |
| --- | --- | --- | --- | --- | --- | --- | --- | --- | --- | --- | --- | --- |
|  |  | Sessions included | 24 | 64 | 76 | 162 | 15 | 159 | 72 | 17 | 167 | 142 |
|  |  | Outlier Sessions excluded | 1 | 10 | 10 | 15 | 0 | 27 | 3 | 1 | 12 | 12 |
| SNR | PD/T2 | Mean | 0.45 | 0.2 | 0.32 | 0.34 | 0.44 | 0.15 | 0.28 | 0.22 | 0.58 | 0.08 |
|  |  | StdDev | 0.37 | 0.26 | 0.26 | 0.26 | 0.34 | 0.18 | 0.27 | 0.3 | 0.2 | 0.12 |
|  | 3DT1 | Mean | 0.48 | 0.41 | 0.48 | 0.5 | 0.15 | 0.4 | 0.49 | 0.44 | 0.27 | 0.33 |
|  |  | StdDev | 0.28 | 0.22 | 0.25 | 0.19 | 0.1 | 0.21 | 0.22 | 0.29 | 0.1 | 0.19 |
|  | T2-star | Mean | 0.6 | 0.42 | 0.13 | 0.17 | 0.7 | 0.45 | 0.48 | 0.42 | 0.07 | 0.46 |
|  |  | StdDev | 0.23 | 0.2 | 0.32 | 0.29 | 0.29 | 0.27 | 0.31 | 0.26 | 0.07 | 0.21 |
|  | 2D Flair | Mean | 0.59 | 0.52 | 0.5 | 0.54 | 0.44 | 0.48 | 0.46 | 0.57 | 0.52 | 0.38 |
|  |  | StdDev | 0.28 | 0.22 | 0.26 | 0.18 | 0.32 | 0.18 | 0.19 | 0.26 | 0.18 | 0.26 |
| CNR | PD/T2 | Mean | 0.25 | 0.16 | 0.24 | 0.32 | 0.33 | 0.12 | 0.29 | 0.27 | 0.3 | 0.1 |
|  |  | StdDev | 0.27 | 0.21 | 0.24 | 0.26 | 0.35 | 0.19 | 0.28 | 0.39 | 0.25 | 0.17 |
|  | 3DT1 | Mean | 0.48 | 0.27 | 0.36 | 0.27 | 0.3 | 0.31 | 0.15 | 0.51 | 0.07 | 0.26 |
|  |  | StdDev | 0.33 | 0.21 | 0.27 | 0.22 | 0.3 | 0.22 | 0.13 | 0.29 | 0.06 | 0.24 |
|  | T2-star | Mean | 0.28 | 0.26 | 0.09 | 0.24 | 0.41 | 0.28 | 0.21 | 0.3 | 0.29 | 0.27 |
|  |  | StdDev | 0.29 | 0.2 | 0.25 | 0.24 | 0.38 | 0.22 | 0.22 | 0.25 | 0.23 | 0.22 |
|  | 2D Flair | Mean | 0.35 | 0.39 | 0.4 | 0.22 | 0.51 | 0.37 | 0.32 | 0.51 | 0.19 | 0.25 |
|  |  | StdDev | 0.3 | 0.25 | 0.24 | 0.2 | 0.34 | 0.24 | 0.22 | 0.29 | 0.14 | 0.23 |

The cross-sectional view of the Spotfire Dashboard displays these summary statistical measures per site, and the longitudinal view displays QC values over time per site as seen in the left and right plots, respectively, of the middle panel of Figure 4. The results of this pipeline, in conjunction with manual QC, are used as a valuable resource to assess scan quality at each ONDRI site.

### 3.4 Pipeline for monitoring and correction of MR scanner geometric gradient field distortions with the Lego® phantom

Estimated geometric distortions are unique to the individual scanners as shown in Figure 5 for two different ONDRI imaging sites. The mean image distortion (d-_m_) was calculated for all points within a spherical region of radius 100 mm positioned at the magnet isocentre. The average image distortion (d-_m_) measured across all ten ONDRI imaging centres was between 0.43 – 1.38 mm. We have found that the average image distortion is very stable over time. A detailed analysis of the gradient field distortions at each MRI site, the method used for correction, and characterization of the temporal variance of the distortion will be presented in a follow-up manuscript.

**Figure 5:**
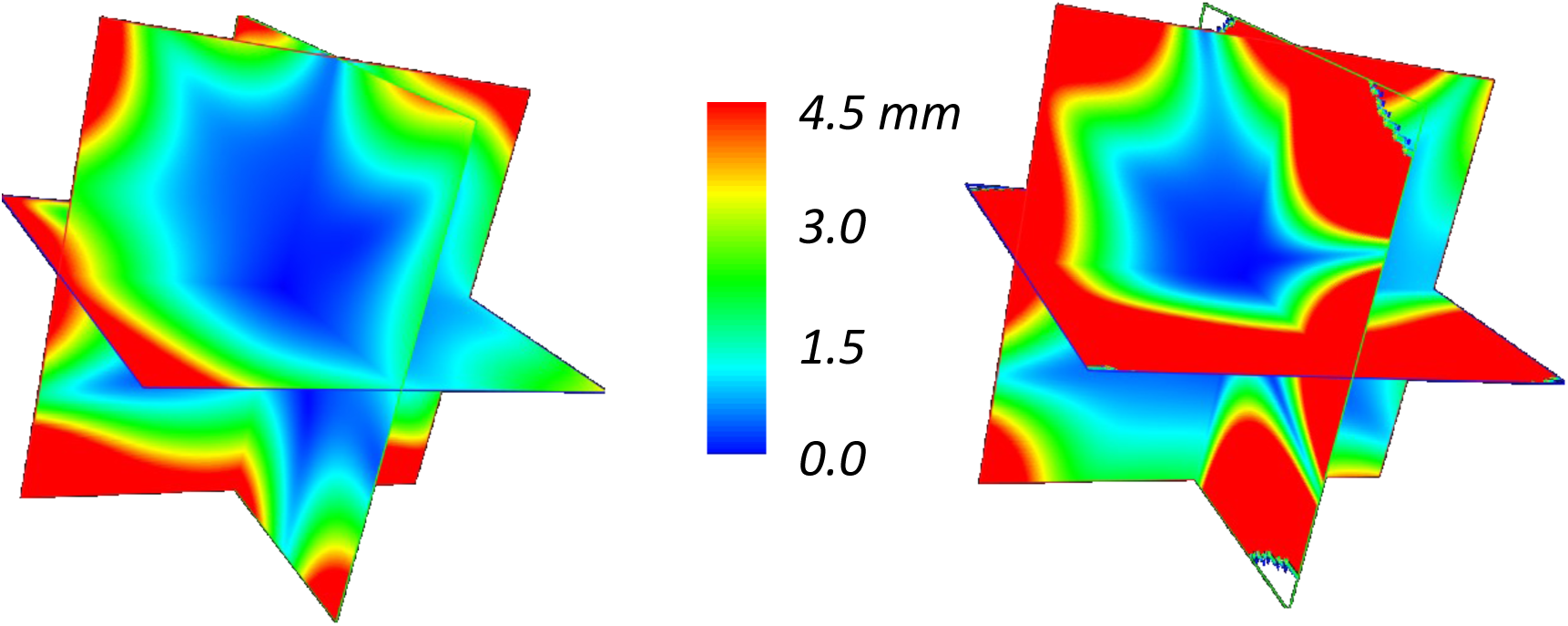
Estimated geometric distortions of a 30 × 30 × 30 cm volume centered at isocentre for two different ONDRI centers. Each scanner has a unique distortion characteristic when estimated using the LEGO® phantom.

### 3.5 Automatic DTI quality assessment pipeline

To perform phantom scans, scan parameters were matched to the human participant protocol and the phantom was scanned twice: with and without parallel imaging (PAR and NPAR respectively). All metrics except Nyquist ghosting (only available for NPAR scans) were then computed twice for both sets of scans.

Many intermediate figures are produced to track, troubleshoot, and verify the results. The following cumulative metrics were extracted from the DTI-QA tool: average (AVE) FA, standard deviation (STD) FA, AVE column pixel shift, AVE SNR of B0s, AVE SNR of non-DWIs, coefficient of variation of SNR across DWIs, AVE Nyquist ghost ratio, and AVE B0 distortion (Figures 6 and 7). These metrics are tabulated and plotted on a web-based dashboard created using Spotfire, accessible to multisite investigators in BrainCODE. Cross-sectional and longitudinal views track the stability of results across and within sites, respectively; the outliers are then identified and tracked.

**Figure 6.**
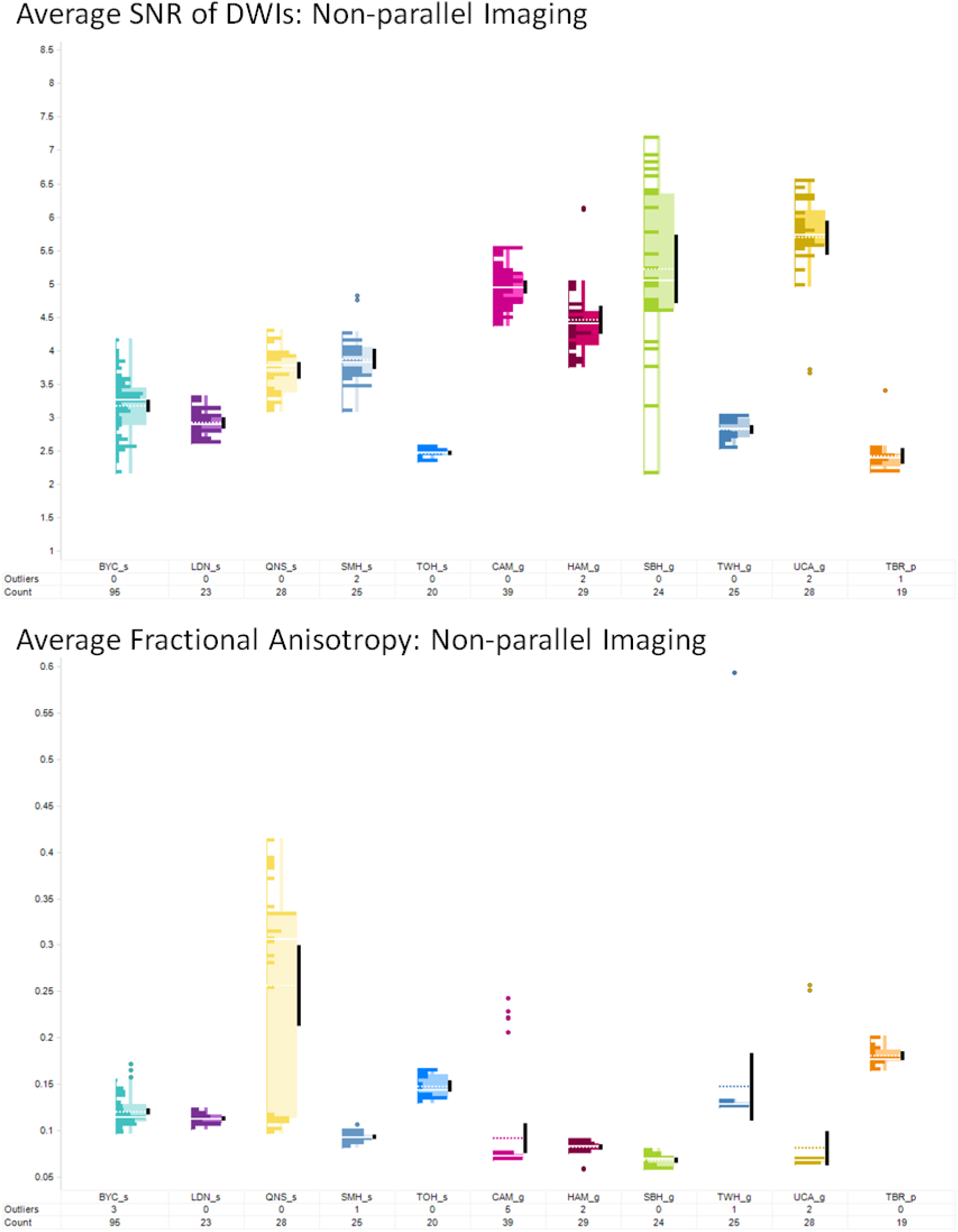
Assessment of the DTI imaging across sites.

**Figure 7.**
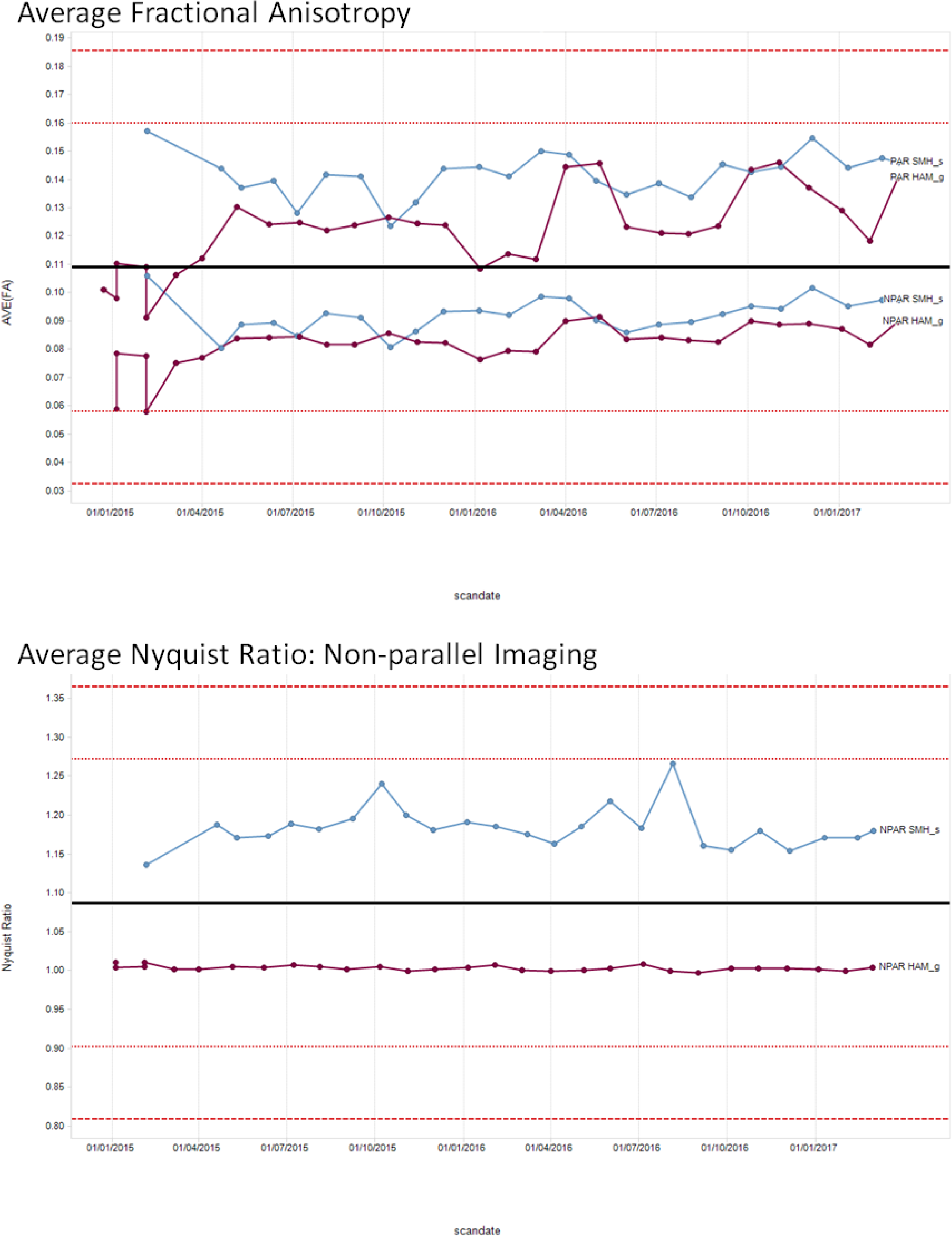
Example assessment of DTI metrics over time to assess stability.

### 3.6 Automatic Resting State fMRI scanner performance monitoring pipelines (fBIRN)

Table 4 displays the aggregated results of the fBIRN pipeline. On average, outlier data (i.e., data that exceed +/-3.00 standard deviations from a given site’s historical mean) account for 5.5% (SD = 3.7%) of each site’s data upload.

**Table 4.** fBIRN pipeline results

|  |  | BYC | CAM | HAM | LDN | QNS | SBH | SMH | TBR | TOH | TWH |
| --- | --- | --- | --- | --- | --- | --- | --- | --- | --- | --- | --- |
|  | Count | 122 | 37 | 31 | 27 | 29 | 27 | 25 | 21 | 20 | 26 |
|  | Excluded* | 0 | 11 | 0 | 0 | 7 | 0 | 2 | 1 | 1 | 4 |
|  | Outliers | 6 | 2 | 1 | 1 | 1 | 0 | 2 | 2 | 2 | 0 |
| Mean Signal | Mean | 1587.07 | 5914.78 | 6325.00 | 1421.40 | 1474.89 | 6953.50 | 1149.15 | 1703.25 | 1310.72 | 3581.47 |
|  | StdDev | 93.65 | 176.09 | 1230.95 | 111.55 | 55.56 | 718.41 | 35.45 | 117.63 | 36.52 | 219.71 |
| SNR | Mean | 167.00 | 312.48 | 250.30 | 194.49 | 182.08 | 489.67 | 207.96 | 254.88 | 166.87 | 324.70 |
|  | StdDev | 12.48 | 20.42 | 15.07 | 13.95 | 18.14 | 49.33 | 13.18 | 18.92 | 8.70 | 23.23 |
| SFNR | Mean | 167.13 | 309.56 | 248.79 | 197.01 | 180.85 | 448.14 | 206.79 | 236.40 | 167.78 | 317.58 |
|  | StdDev | 9.69 | 11.39 | 11.91 | 9.83 | 12.70 | 34.71 | 9.30 | 6.33 | 4.00 | 10.22 |
| Std. Dev. | Mean | 1.25 | 2.68 | 4.11 | 0.68 | 1.16 | 5.29 | 0.62 | 2.79 | 1.05 | 2.98 |
|  | StdDev | 0.15 | 1.03 | 1.56 | 0.07 | 0.10 | 2.52 | 0.47 | 0.36 | 0.11 | 0.32 |
| Percent Fluctuation | Mean | 0.08 | 0.04 | 0.06 | 0.05 | 0.08 | 0.07 | 0.05 | 0.16 | 0.08 | 0.08 |
|  | StdDev | 0.01 | 0.02 | 0.02 | 0.00 | 0.01 | 0.03 | 0.04 | 0.02 | 0.01 | 0.01 |
| Drift | Mean | 0.75 | 0.48 | 1.26 | 0.94 | 0.91 | 0.69 | 0.49 | 1.42 | 0.64 | 2.17 |
|  | StdDev | 0.17 | 0.22 | 0.35 | 0.2 | 0.26 | 0.18 | 0.24 | 0.34 | 0.15 | 0.27 |
| Drift Fit | Mean | 0.39 | 0.29 | 1.02 | 0.72 | 0.51 | 0.37 | 0.27 | 0.82 | 0.29 | 1.90 |
|  | StdDev | 0.18 | 0.16 | 0.35 | 0.19 | 0.24 | 0.14 | 0.17 | 0.32 | 0.15 | 0.26 |
| Mean Ghost | Mean | 1.67 | 0.38 | 0.39 | 1.64 | 3.07 | 2.43 | 2.93 | 0.89 | 2.72 | 0.32 |
|  | StdDev | 0.09 | 0.04 | 0.11 | 0.18 | 0.74 | 0.37 | 0.10 | 0.34 | 0.08 | 0.03 |
| Mean Bright Ghost | Mean | 3.37 | 2.12 | 2.12 | 2.98 | 8.26 | 8.06 | 5.87 | 3.41 | 6.01 | 2.30 |
|  | StdDev | 0.19 | 0.21 | 0.44 | 0.27 | 4.47 | 1.80 | 0.24 | 0.85 | 0.46 | 0.18 |
| Max FWHMX | Mean | 2.73 | 4.33 | 4.24 | 2.91 | 2.77 | 5.46 | 2.94 | 5.07 | 2.74 | 4.46 |
|  | StdDev | 0.09 | 0.32 | 0.33 | 0.02 | 0.08 | 1.39 | 0.78 | 0.48 | 0.18 | 0.12 |
| Max FWHMY | Mean | 2.93 | 3.98 | 3.95 | 3.08 | 2.81 | 5.05 | 3.22 | 5.13 | 3.04 | 4.30 |
|  | StdDev | 0.09 | 0.24 | 0.29 | 0.02 | 0.29 | 1.30 | 0.76 | 0.62 | 0.16 | 0.13 |
| Max FWHMZ | Mean | 1.92 | 2.71 | 2.45 | 1.97 | 2.07 | 3.39 | 2.27 | 3.07 | 2.06 | 2.66 |
|  | StdDev | 0.06 | 0.17 | 0.35 | 0.05 | 0.18 | 0.69 | 0.55 | 0.74 | 0.16 | 0.14 |
| Min FWHMX | Mean | 2.54 | 3.85 | 3.89 | 2.77 | 2.53 | 4.25 | 2.50 | 4.22 | 2.45 | 3.97 |
|  | StdDev | 0.02 | 0.15 | 0.15 | 0.02 | 0.06 | 0.31 | 0.03 | 0.07 | 0.01 | 0.03 |
| Min FWHMY | Mean | 2.77 | 3.56 | 3.60 | 2.95 | 2.51 | 3.84 | 2.86 | 4.21 | 2.82 | 3.89 |
|  | StdDev | 0.01 | 0.12 | 0.12 | 0.02 | 0.46 | 0.31 | 0.03 | 0.07 | 0.01 | 0.05 |
| Min FWHMZ | Mean | 1.31 | 1.32 | 1.30 | 1.26 | 1.37 | 1.47 | 1.30 | 1.41 | 1.31 | 1.34 |
|  | StdDev | 0.08 | 0.08 | 0.10 | 0.07 | 0.19 | 0.15 | 0.08 | 0.09 | 0.06 | 0.10 |
| RDC | Mean | 7.42 | 7.66 | 6.71 | 11.13 | 7.15 | 3.59 | 11.02 | 2.69 | 7.63 | 3.94 |
|  | StdDev | 0.93 | 1.13 | 1.22 | 0.83 | 1.42 | 1.43 | 3.08 | 0.25 | 0.80 | 0.39 |

### 3.7 Manual procedures for image assessment

Of the 989 scans acquired up until the ‘new enrollment’ cut-off date, manual visual review of the imaging was able to identify 22 sessions that were unusable, based on the quality of the T1-weighted scan. Of the 22 failures, 15 were successfully able to be reacquired, while 7 were unable to be reacquired (e.g., patient refused, excessive motion in all sequences so unlikely to improve, mobility issues). Further assessment of the remaining sequences show higher rates of individual sequence failure but still allow for some overall usability of the data.

## 4. DISCUSSION

The neuroinformatics procedures and pipelines employed in ONDRI attempt to deal with many of the challenges associated with generating MR imaging data from large multi-site studies. As with many other studies, some basic measures were taken to account for known issues, such as visual inspection for motion and other artifacts, and various types of distortion and inhomogeneity correction in the images. Similarly, considerable effort was put into harmonizing the image acquisition protocols as a critical step towards generating comparable data between different scanner makes and models. However, where ONDRI surpasses many other studies is with the thoroughness and attention to detail at multiple levels, from naming and protocol adherence, to sequence specific SNR/CNR measures and phantom imaging tracking scanner performance, in order to insure that the data are of the highest possible caliber.

The ONDRI study used two separate phantoms to monitor scanner stability and correct gradient field non-linear distortions. This increased the complexity and time associated with obtaining various measures (both in scan time and post processing) compared to other studies that use a single phantom. However, it was a very cost effective alternative that took advantage of freely available fBIRN tools and pipelines to obtain detailed quality control data to monitor scanner performance over time.

Our criteria for determining the overall pass or fail of any given imaging session was based solely on the quality of the T1-weighted image. This was chosen as the basis for overall usability due to the fact that many of the primary outcome measures could still be generated from the data, and because the T1 served as the foundation for the usability of many other sequence types, such as the resting state fMRI and DTI. Without the T1, very few of the other sequences could generate meaningful data, while with the T1 (and potentially without the other sequences), considerable measures could still be obtained.

Early on, ensuring that sites properly adhered to the imaging protocol and output correctly named images was the greatest challenge, however, the primary cause for this was human error. Once sites became familiar with these procedures, the protocol and naming adherence issues were of less concern (although they never fully went away). Another source of human error at the outset of the project, and a great challenge to the overall ability to perform timely quality control was with the delay between data acquisition and upload to the central repository. While sites were instructed to upload the data within 48 hours of acquisition, some sites were rather delinquent and scans were routinely uploaded weeks and occasionally months after they were acquired. This delay-to-upload improved over the course of the project with greater enforcement and consistent reminders to the sites and coordinators.

Protocol adherence was also a greater challenge for Siemens scanners than was initially anticipated, as the more modern scanners tended to make frequent user-independent on-the-fly modifications to parameters such as TR and TE, to automatically adjust for SAR, which often pushed these values outside of the expected ranges. However, through manual review of each of the protocol deviations, we were able to determine if they were still acceptable and within tolerances that would allow for meaningful and comparable data to be generated.

The quality assurance and quality control measures implemented for the acquisition of magnetic resonance imaging data as part of the multi-site Ontario neurodegenerative disease research initiative have produced a robust database of images that span several neurodegenerative diseases and now allow careful comparison between cohorts and over time. Using rigorous approaches to standardization and monitoring of compliance we have achieved high quality useable imaging data in more than 99% of participants. Continuous monitoring and quality control is required to ensure compliance of all aspects related to data acquisition and processing.

