## Supplementary_Table_1 for "An overview of the quality assurance and quality control of magnetic resonance imaging data for the Ontario Neurodegenerative Disease Research Initiative (ONDRI): pipeline development and neuroinformatics"

### Supplementary Table 1 - MRI Acquisition Protocols

| STUDY | OBI - ONDRI | | |
|---|---|---|---|
| SEQUENCE | 3DT1 | | |
| **Protocol** | | | |
| Vendor | GE | Philips | Siemens |
| Field Strength | 3T | 3T | 3T |
| Model | Discovery | Achieva | Skyra/Trio/Prisma |
| Version | 22 | 3.2.3 | |
| Sequence Name | 3D FAST SPGR | 3D TFE | 3D MP-RAGE |
| Imaging Options | IrP- Asset | Fast (Sense) | iPat |
| **Pulse Timing** | | | |
| TE (ms) | Min full | Min (3.3) | 2.98 |
| TR (ms) | Min | Min (7.3) | 2300 |
| Flip Angle (°) | 11 | 9 | 9 |
| TI (ms) | 400 | 945 | 900 |
| **Scan Range** | | | |
| FOV (in-plane) (mm) | 256 x 256 | 256 x 248 | 256 x 256 |
| Slice Thickness (mm) | 1 | 1 | 1 |
| Gap Between Slices (mm) | 0 | 0 | 0 |
| No. Slices | 176 | 176 | 176 |
| **Acquisition** | | | |
| Orientation | Sagittal | Sagittal | Sagittal |
| Matrix Size | 256 x 256 | 256 x 248 | 256 x 256 |
| Voxel Size [L/R x A/P x I/S] | 1 x 1 x 1 | 1 x 1 x 1 | 1 x 1 x 1 |
| NEX | 1 | 1 | 1 |
| Acceleration Factor (Parallel factor*) | 2 | 2 | 2 |
| **Other** | | | |
| Fat Suppression | None | None | None |
| Bandwidth | 31.25 (kHz) | 228 (Hz/px) | 240 (Hz/px) |
| Echo Train Length | - | - | - |
| **Coil Type** | | | |
| Head | X | X | X |
| Channel | 8-12 (HNS) | 8 | 12(Trio) 20 (Prisma) |
| **Time** | | | |
| PRESCAN TIME+ | 00:30 | 00:30 | 00:30 |
| SCAN TIME | 04:52 | 06:17 | 05:21 |
| TOTAL TIME (MIN) | 05:22 | 06:47 | 05:51 |

| | | | |
|---|---|---|---|
| ***Study*** | ***OBI - ONDRI*** | | |
| ***Sequence*** | ***PD/T2*** | | |
| ***Protocol*** | | | |
| Vendor | GE | Philips | Siemens |
| Field Strength | 3T | 3T | 3T |
| Model | Discovery | Achieva | Skyra/Trio/Prisma |
| Version | 22 | 3.2.3 | |
| Sequence Name | FSE-XL | TSE | TSE |
| Imaging Options | EDR, Asset | Fast (Sense) | iPat |
| ***Pulse Timing*** | | | |
| TE (ms) (2 echo scan) | Min full/86 | 13/100 | 10/93 |
| TR (ms) | 3000 | 3000 | 3000 |
| Flip Angle (°) | 125 | 90 | 165 |
| TI (ms) | - | - | - |
| ***Scan Range*** | | | |
| FOV (in-plane) (mm) | 240 x 240 | 240 x 240 | 240 x 240 |
| Phase FOV | 75% | 75% | 81% |
| Slice Thickness (mm) | 3 | 3 | 3 |
| Gap Between Slices (mm) | 0 | 0 | 0 |
| No. Slices | 48 | 48 | 48 |
| ***Acquisition*** | | | |
| Orientation | Oblique Axial | Oblique Axial | Oblique Axial |
| Matrix Size | 256 x 256 | 256 x 254 | 256 x 256 |
| Voxel Size [L/R x A/P x I/S] | 0.94 x 0.94 x 3 | 0.94x 0.94 x 3 | 0.94 x 0.94 x 3 |
| NEX | 1 | 1 | 1 |
| Acceleration Factor (Parallel factor*) | 2 | 2 | 2 |
| ***Other*** | | | |
| Fat Suppression | Yes (FAT-SAT) | Yes | Yes |
| Bandwidth | 20 (kHz) | 222 (Hz/px) | 181 (Hz/px) |
| Echo Train Length | 12 | 12 | 14 |
| ***Coil Type*** | | | |
| Head | X | X | X |
| Channel | 8-12 (HNS) | 8 | 12(Trio) 20 (Prisma) |
| ***Time*** | | | |
| ***Prescan Time+*** | 00:30 | 00:30 | 00:30 |
| ***Scan Time*** | 02:43 | 04:12 | 03:11 |
| ***Total Time (min)*** | 03:13 | 04:42 | 03:41 |

| | | | |
|---|---|---|---|
| ***Study*** | ***OBI - ONDRI*** | | |
| ***Sequence*** | ***2D FLAIR*** | | |
| ***Protocol*** | | | |
| Vendor | GE | Philips | Siemens |
| Field Strength | 3T | 3T | 3T |
| Model | Discovery | Achieva | Skyra/Trio/Prisma |
| Version | 22 | 3.2.3 | |
| Sequence Name | 2D T2FLAIR | 2D IR TSE | 2D IR TDF |
| Imaging Options | EDR, IR | Fast (Sense) | iPat |
| ***Pulse Timing*** | | | |
| TE (ms) | 140 | 125 | 120 |
| TR (ms) | 9000 | 9000 | 9000 |
| Flip Angle (°) | 125 | 90 (150 refocus) | 165 |
| TI (ms) | 2250 | 2500 | 2500 |
| ***Scan Range*** | | | |
| FOV (in-plane) (mm) | 240 x 240 | 240 x 240 | 240 x 240 |
| Slice Thickness (mm) | 3 | 3 | 3 |
| Gap Between Slices (mm) | 0 | 0 | 0 |
| No. Slices | 48 | 48 | 48 |
| ***Acquisition*** | | | |
| Orientation | Oblique Axial | Oblique Axial | Oblique Axial |
| Matrix Size | 256 x 256 | 256 x 242 | 256 x 256 |
| Voxel Size [L/R x A/P x I/S] | 0.94 x 0.94 x 3 | 0.94 x 0.99 x 3 | 0.94 x 0.94 x 3 |
| NEX | 1 | 1 | 1 |
| Acceleration Factor (Parallel factor*) | No Asset | 2 (SENSE) | 2 |
| ***Other*** | | | |
| Fat Suppression | None | None | None |
| Bandwidth | 25 (kHz) | 242 (Hz/px) | 220 (Hz/px) |
| Echo Train Length | | 19 | 19 |
| ***Coil Type*** | | | |
| Head | X | X | X |
| Channel | 8-12 (HNS) | 8 | 12(Trio) 20 (Prisma) |
| ***Time*** | | | |
| ***Prescan Time+*** | 00:30 | 00:30 | 00:30 |
| ***Scan Time*** | 04:32 | 03:45 | 02:44 |
| ***Total Time (min)*** | 05:02 | 04:15 | 03:14 |

| | | | |
|---|---|---|---|
| ***STUDY*** | ***OBI - ONDRI*** | | |
| ***SEQUENCE*** | ***T2-star*** | | |
| ***Protocol*** | | | |
| Vendor | GE | Philips | Siemens |
| Field Strength | 3T | 3T | 3T |
| Model | Discovery | Achieva | Skyra/Trio/Prisma |
| Version | 22 | 3.2.3 | |
| Sequence Name | GRE | FFE | GRE |
| Imaging Options | - | Sense | iPat |
| ***Pulse Timing*** | | | |
| TE (ms) | 20 | 21 | 20 |
| TR (ms) | 650 | 650 | 650 |
| Flip Angle (°) | 20 | 20 | 20 |
| TI (ms) | - | - | - |
| ***Scan Range*** | | | |
| FOV (in-plane) (mm) | 240 x 240 | 240 x 240 | 240 x 240 |
| Phase FOV | 75% | 75% | 75% |
| Slice Thickness (mm) | 3 | 3 | 3 |
| Gap Between Slices (mm) | 0 | 0 | 0 |
| No. Slices | 48 | 48 | 48 |
| ***Acquisition*** | | | |
| Orientation | Oblique Axial | Oblique Axial | Oblique Axial |
| Matrix Size | 256 x 256 | 256 x 256 | 256 x 256 |
| Voxel Size [L/R x A/P x I/S] | 0.94 x 0.94 x 3 | 0.94 x 0.94 x 3 | 0.94 x 0.94 x 3 |
| NEX | 1 | 1 | 1 |
| Acceleration Factor (Parallel factor*) | No Asset | 2 (SENSE) | 2 |
| ***Other*** | | | |
| Fat Suppression | None | None | None |
| Bandwidth | 19.23 (kHz) | 217 (Hz/px) | 200 (Hz/px) |
| Echo Train Length | 1 | 1 | 1 |
| CV act_te (GE only) | 20000 | | |
| ***Coil Type*** | | | |
| Head | X | X | X |
| Channel | 8-12 (HNS) | 8 | 12(Trio) 20 (Prisma) |
| ***Time*** | | | |
| ***PRESCAN TIME+*** | 00:30 | 00:30 | 00:30 |
| ***SCAN TIME*** | 02:15 | 02:52 | 03:04 |
| ***TOTAL TIME (MIN)*** | | | |

| **STUDY** | ***OBI - ONDRI*** | | |
|---|---|---|---|
| **SEQUENCE** | ***fMRI-RS*** | | |
| ***Protocol*** | | | |
| Vendor | GE | Philips | Siemens |
| Field Strength | 3T | 3T | 3T |
| Model | Discovery | Achieva | Skyra/Trio/Prisma |
| Version | 22 | 3.2.3 | |
| Sequence Name | fMRI EPI | fMRI EPI | fMRI EPI |
| Imaging Options | EDR, “eyes open” | GRE EPI, CLEAR SENSE, “eyes open” | “eyes open” |
| ***Pulse Timing*** | | | |
| TE (ms) | 30 | 30 | 30 |
| TR (ms) | 2400 | 2400 | 2400 |
| Flip Angle (°) | 70 | 70 | 70 |
| TI (ms) | - | - | - |
| ***Scan Range*** | | | |
| FOV (in-plane) (mm) | 224 x224 | 224 x 224 | 224 x 224 |
| Slice Thickness (mm) | 3.5 | 3.5 | 3.5 |
| Gap Between Slices (mm) | 0 | 0 | 0 |
| No. Slices | 41 | 41 | 41 |
| ***Acquisition*** | | | |
| Orientation | Oblique Axial | Oblique Axial | Oblique Axial |
| Matrix Size | 64 x 64 | 64 x 64 | 64 x 64 |
| Voxel Size [L/R x A/P x I/S] | 3.5 x 3.5 x 3.5 | 3.5 x 3.5 x 3.5 | 3.5 x 3.5 x 3.5 |
| NEX | 1 | 1 | 1 |
| Acceleration Factor (Parallel factor*) | 2 | 2 | 2 |
| Slice Order | | Ascending | |
| ***Other*** | | | |
| Fat Suppression | Fat Sat. | Fat Sat. SPIR | Fat Sat. |
| Bandwidth (Hz/Px) | 7812 | 2441 | 2440 |
| Number of acquisitions | 250 | 250 | 250 |
| EPI Factor | | 35 | 31 |
| ***Coil Type*** | | | |
| Head | X | X | X |
| Channel | 8-12 (HNS) | 8 | 12(Trio) 20 (Prisma) |
| ***Time*** | | | |
| ***PRESCAN TIME+*** | 00:30 | 00:30 | 00:30 |
| ***SCAN TIME*** | 10:00 | 10:00 | 10:00 |
| ***TOTAL TIME (MIN)*** | 10:30 | 10:30 | 10:30 |

| | | | | |
|---|---|---|---|---|
| ***Study*** | ***OBI - ONDRI*** | | | |
| ***Sequence*** | ***DTI*** | | | |
| ***Protocol*** | | | | |
| Vendor | GE | Philips | Siemens | Siemens |
| Field Strength | 3T | 3T | 3T | 3T |
| Model | Discovery | Achieva | Skyra/Trio/Prisma | Skyra/Trio/Prisma |
| Version | 22 | 3.2.3 | | 17 |
| Sequence Name | DWI | DWI | DWI | DWI |
| Imaging Options | ASSET | Sense | iPat | iPat |
| ***Pulse Timing*** | | | | |
| TE (ms) | Min | 100 | 96/ 63(Prisma) | 96/ 64 (Prisma) |
| TR (ms) | 9000 | 9931 | 9400 | 9400 |
| Flip Angle (°) | 90 | 90 | 90 | 90 |
| TI (ms) | - | - | - | - |
| ***Scan Range*** | | | | |
| FOV (in-plane) (mm) | 256 x 256 | 256x 256 | 256 x 256 | 256 x 256 |
| Slice Thickness (mm) | 2 | 2 | 2 | 2 |
| Gap Between slices (mm) | 0 | 0 | 0 | 0 |
| No. Slices | 70 | 70 | 70 | 70 |
| ***Acquisition*** | | | | |
| Orientation | Oblique Axial | Oblique Axial | Oblique Axial | Oblique Axial |
| Matrix Size | 128 x 128 | 128 x 128 | 128 x 128 | 128 x 128 |
| Voxel Size (L/R x A/P x I/S) | 2 x 2 x 2 | 2 x 2 x 2 | 2 x 2 x 2 | 2 x 2 x 2 |
| NEX | 1 | 1 | 1 | 1 |
| Acceleration Factor (Parallel factor*) | 2 | 2 | 2 | 2 |
| ***Diffusion*** | | | | |
| b-value 1 | 0 | 0 | 0 | 0 |
| b-value 2 | 1000 | 1000 | 1000 | |
| Number of Directions | 30 | 32 | 30 | 1 |
| ***Other*** | | | | |
| Fat Suppression | FatSat | FatSat | FatSat | FatSat |
| Bandwidth (Hz/Px) | | 2045 | 2056 | 2056 |
| T2 Images | 3 | 3 | 3 | |
| EPI Factor | | 67 | 128 | 128 |
| Gradients | | | Monopolar (Prisma) | Monopolar (Prisma) |
| CV rhimsize (GE only) | 128 | | | |
| ***Coil Type*** | | | | |

| | | | | |
|---|---|---|---|---|
| Head | x | x | x | x |
| Channel | 8-12 (HNS) | 8 | 12(Trio) 20 (Prisma) | 12(Trio) 20 (Prisma) |
| ***Time*** | | | | |
| **PRESCAN TIME+** | 00:30 | 00:30 | 00:30 | 00:30 |
| **SCAN TIME** | 05:06 | 0:6:58 | 05:21 | 00:39 |
| **TOTAL TIME (MIN)** | 05:36 | 07:28 | 05:33 | 01:08 |

#### Core Protocol Scan Time Estimates

| | *GE* | *Philips* | *Siemens* |
|---|---|---|---|
| Set up | *05:00* | *05:00* | *05:00* |
| 3DT1 | 05:22 | 06:47 | 05:51 |
| PD/T2 | 02:49 | 04:42 | 03:41 |
| 2D FLAIR | 05:02 | 04:15 | 03:14 |
| T2-star *or* T2-star_PM *or* T2-star_PMRI | 03:45 | 03:22 | 03:34 |
| RS-fMRI | 10:30 | 10:30 | 10:30 |
| DTI | 06:30 | 07:28 | 06:41 |
| Set down | *02:30* | *02:30* | *02:30* |
| **Total scan time** | 33:58 | 37:04 | 33:31 |
| **Total session time** | **41:28** | **44:34** | **41:01** |
